# COMPOSITUM 1 (COM1) contributes to the architectural simplification of barley inflorescence via meristem identity signals

**DOI:** 10.1101/2020.02.18.952705

**Authors:** N. Poursarebani, C. Trautewig, M. Melzer, T. Nussbaumer, U. Lundqvist, T. Rutten, T. Schmutzer, R. Brandt, A. Himmelbach, L. Altschmied, R. Koppolu, H. M. Youssef, R. Sibout, M. Dalmais, A. Bendahmane, N. Stein, Z. Xin, T. Schnurbusch

**Author notes:** Corresponding authors. (NP); (TS).

## Abstract

Grasses have varying inflorescence shapes; however, little is known about the genetic mechanisms specifying such shapes among tribes. We identified the grass-specific TCP transcription factor COMPOSITUM 1 (COM1) expressed in inflorescence meristematic boundaries of different grasses. COM1 specifies branch-inhibition in Triticeae (barley) versus branch-formation in non-Triticeae grasses. Analyses of cell size, cell walls and transcripts revealed barley COM1 regulates cell growth, affecting cell wall properties and signaling specifically in meristematic boundaries to establish identity of adjacent meristems. *COM1* acts upstream of the boundary gene *Liguleless1* and confers meristem identity partially independent of the *COM2* pathway. Furthermore, COM1 is subject to purifying natural selection, thereby contributing to specification of the spike inflorescence shape. This meristem identity pathway has conceptual implications for both inflorescence evolution and molecular breeding in Triticeae.

## Introduction

The grass family (Poaceae), one of the largest angiosperm families, has evolved a striking diversity of inflorescence morphologies bearing complex structures such as branches and specialized spikelets ^1^. These structural features are key for sorting the grass family into tribes ^1^. Current grass inflorescences are proposed to originate from a primitive ancestral shape exhibiting “a relatively small panicle-like branching system made up of primary and secondary paracladia (branches), each one standing single at the nodes” ^2^ (**Fig. 1A**). This ancestral panicle-like inflorescence is also known as a compound spike ^3–5^. Several independent or combined diversification processes throughout the evolutionary history of the grass family have resulted in the broad diversity of today’s grass inflorescences ^2,3,6^. Some tribes, e.g. Oryzeae (rice) and Andropogoneae (maize and sorghum), still display ancestral and complex compound shapes, keeping true-lateral long primary and secondary branches. Other grasses, such as *Brachypodium distachyon*, show lower inflorescence complexity with branch length and number reduced to lateral, small pedicels ending in only one multi-floretted spikelet (**Fig. 1A-C**). Inflorescences within the tribe Triticeae, e.g. barley (*Hordeum vulgare* L.), probably evolved from the ancestral compound spike into the typical unbranched spike (**Fig. 1D)**. The spike displays the least-complex inflorescence shape due to the sessile nature of spikelets and reduction in rachis internodes ^2,7^. Architectural variation is often manifested through subtle modifications of transcriptional programs during critical transitional windows of inflorescence meristem (IM) maturation ^7,8^ or functional divergence of key transcriptional regulators and/or other genes ^9,10^. Identification of key genetic determinants is crucial for better understanding and explaining both the origin of grass inflorescence diversity and grass developmental gene evolution. Inflorescence developmental patterning controls pollination, grain set and grain number, and is thus highly relevant to agronomy as a target of natural and human selection.

**Fig. 1.**
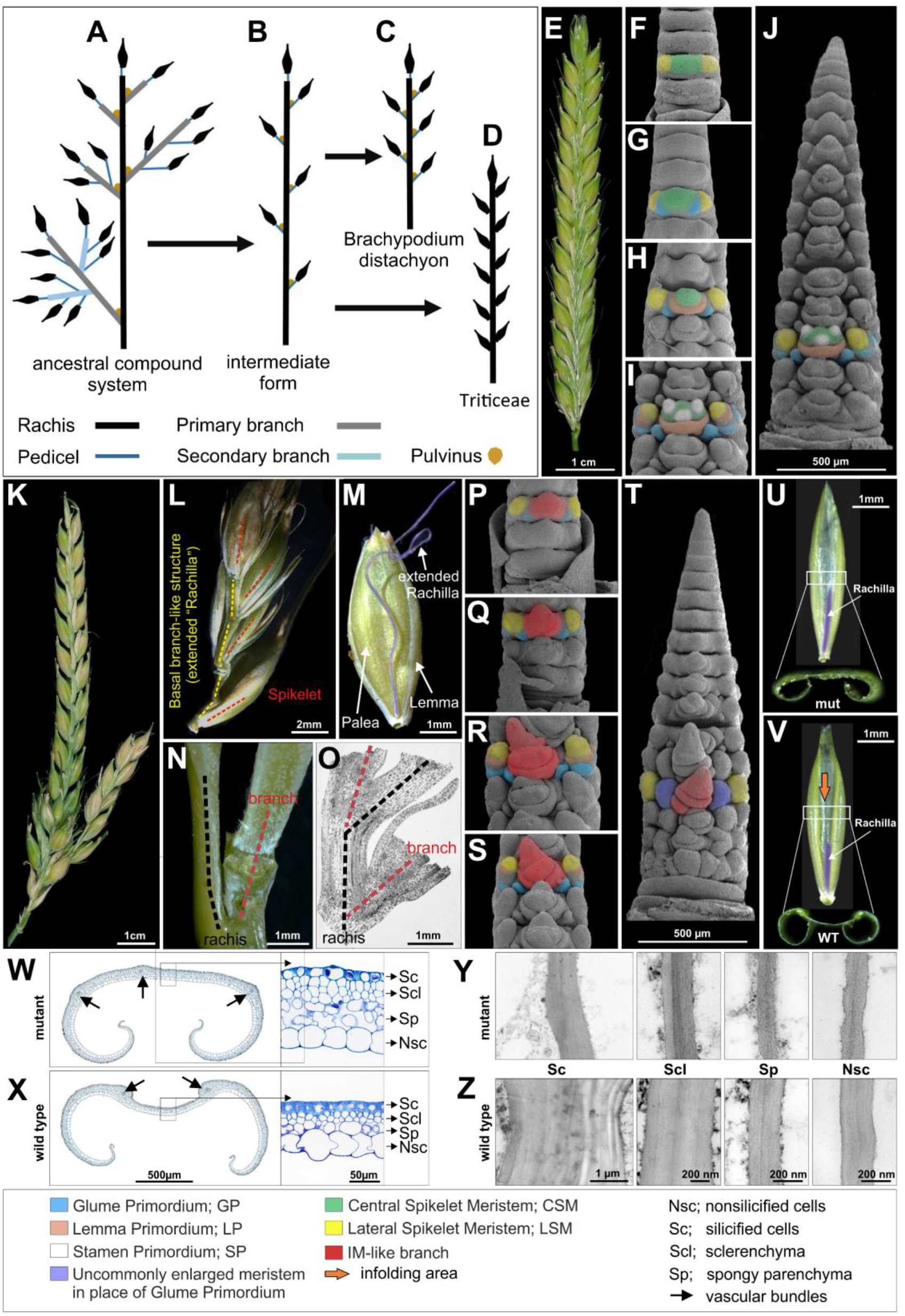
Proposed evolutionary pattern of grass inflorescences, and the spike/palea morphology of wild-type and *com1.a* mutant e.g. BW-NIL(*com1.a*) barley. (**A-D**) Model for grass inflorescence evolution from ancestral compound form to spike in Triticeae; re-drawn from ^2^. (**E**) Spike morphology of wild-type (Wt), two-rowed barley *cv.* Bowman. **(F-I**) SEM imagining of the early developmental stages in immature Wt spike; triple mound: TM (F), glume primordium: GP (**G**), lemma primordium: LP (**H**) and stamen primordium: SP (**I**). Images are taken from basal nodes where a single node is used for color coding. (J) Dorsal view of whole immature Wt spike at stamen primordium stage. (K) Branched spike of BW-NIL(*com1.a*) mutant at maturity. (**L-M**) Depicted is a small, spike-like branch structure, arisen from the central spikelet position due to loss of CSM identity, from intense (**L**) to weak appearance as an extended (ext.) rachilla (**M**), M also depicts a developing grain enclosed by lemma and palea. (N-O) Lack of pulvinus at the base of a branch in BW-NIL(*com1.a*) mutant spike (N) supported by histological imaging (O). (**P-S**) Developmental stages of immature BW-NIL(*com1.a*) mutant spike from early GP (**P**), GP (**Q**), to LP (**R**) and early SP (**S**) taken from the basal nodes. (**T**) Dorsal view of whole immature BW-NIL(*com1.a*) mutant spike at early stamen primordia. (**U)** Longitudinal adaxial view of the palea in BW-NIL(*com1.a*); white rectangle corresponds to the area used to take sections for histologica l analysis and to the lower image depicting the flat-plane surface of a palea cross section. (V) Longitudinal adaxial view of the palea in Wt; the lower image corresponds to the infolding surface of a palea cross section. (**W-X**) Histological analyses of transverse sections (from U and V; white rectangles) of the palea in BW-NIL(*com1.a*) (**W**) and Wt **(X**). Paleae are from spikelets shortly before anthesis. (**Y-Z**) TEM based imaging of walls of paleae cells in BW-NIL(*com1.a*) (**Y**) versus Wt (**Z**).

A valuable toolkit to explore such genetic determinants regulating inflorescence patterns in Triticeae is a collection of morphological barley mutants, induced by physical and chemical mutagens ^11^. This collection includes both compositum-barleys displaying branched spikes and their corresponding near isogenic lines (NIL) ^12^. There are eight of such NILs reported ^12^ one of which, NIL *COMPOSITUM2 (COM2),* has been characterized so far. The underlying gene encodes an AP2/ERF transcription factor orthologous to maize *BD1* with a conserved function of branch suppression across grass ^13^. Here, we have conducted a detailed phenotypic inspection of another compositum-barley, NIL *COM1*, also displaying non-canonical, i.e. branched, spike morphology. We identified and characterized the underling boundary forming protein, a grass-specific TCP transcription factor, and present evidence that *COM1* in barley has evolved a function opposite to its orthologous genes in maize and rice, *ZmBAD1/WAB1* and *OsREP1/DBOP*, respectively ^14–17^. We further show that its orthologous proteins also functions oppositely in sorghum and *Brachypodium distachyon*. Unlike in these non-Triticeae grasses, in which branch-formation is promoted, *COM1* inhibits spike-branching most likely by affecting meristematic signaling via changing cell wall properties of meristematic boundaries. We generated a double mutant (DM) of *com1.a/com2.g* and provide evidence that DM plants outperformed both single mutants as well as the control wild type plants in supernumerary spikelet formation, and as a consequence, in grain number per spike. Thus, our findings may spur further interests for grass inflorescence evolution but similarly for improving grain number.

## Results

### Atypical for Triticeae—barley *com1.a* forms a branched inflorescence

Barley (and other Triticeae) wild-type (Wt) spikes are typically unbranched and composed of sessile, single-flowered spikelets arranged in a regular distichous fashion of two opposite rows directly attached to the main inflorescence axis, i.e. rachis (**Fig. 1E**). In a mature barley spike, three spikelets per rachis node are visible. Each spikelet initiates from a single meristematic mound first detectable at the triple mound (TM) stage during early reproductive development (**Fig. 1F**). Thus, the TM corresponds to three spikelets meristems (SMs); one central (CSM) and two lateral (LSM) spikelet meristems. The differentiating primordia are followed by several consecutive meristematic and developmental stages e.g. glume primordium (GP; **Fig. 1G**), lemma primordium (LP; **Fig. 1H**), and stamen primordium (SP; **Fig. 1I-J**).^18^

To provide deeper insights into the genetic basis defining inflorescence architecture in Triticeae, we conducted a detailed phenotypic inspection of a NIL of compositum-barley (*com1.a*) mutant displaying an uncommon branched spike (**Fig. 1K)**. The original *com1.a* branched spike mutant was first discovered after simultaneous mutagenesis using EMS and neutron radiation in *cv.* Foma. It was later backcrossed (BC_6_) to a two-rowed barley *cv.* Bowman (BW) ^12^ to create the aforementioned NIL, the BW-NIL(*com1.a*)(**Supplementary Fig. 1A-B**). Thus, we hereafter refer to the BW-NIL(*com1.a*), as *com1.a* mutant. The inflorescence in this *com1.a* mutant, resembles an ancestral compound spike (**Fig. 1K**), but lacks an organ called pulvinus (**Fig. 1N-O**). In non-Triticeae grass species, the pulvinus is present at the axil of lateral long branches in panicles and compound spikes, defining branch angle extent (**Fig. 1A-C**, in brown). We observed differences in spike shape between BW and *com1.a* during early spike differentiation at the late triple mound (TM) to early glume primordium (GP) stage; the mutant central spikelet meristem (SM) is elongated (**Fig. 1P** versus Wt in **1G**), becoming more apparent during later reproductive stages of late glume primordium (**Fig. 1Q**) onwards (**Fig. 1R-S**). At LP, predominantly in the basal part of the spike, meristems of the central spikelet positions undergo evidently SM identity loss, displaying branch- or IM-like meristems (**Fig. 1R**). Instead of generating florets, the meristem continues to elongate and rather functions as an indeterminate spikelet multimer in the form of a primary branch-like structure (**Fig. 1T**). Such branch-like structures occasionally replace other spikelet-related organs, such as the rachilla primordium (RP, the spikelet axis, in **Fig. 1S**; the possible origin for extended rachilla visible at maturity; **Fig. 1L-M**) or glumes (**Fig. 1T** in purple). The *com1* branching phenotype resembles that of the previously described *com2* ^*13*^, in which the formation of branch-like structures results from lack of SM identity (See below).

### *COM1* restricts palea cell size by thickening their cell walls

In barley, the grains are enclosed by two bract-like organs, i.e. lemma and palea, which are part of the floret and provide protection to the developing grain. (Fig. 1M). Besides the branch phenotype, *com1.a* exhibits a deviation in adaxial palea morphology, having a flat plane (**Fig. 1U**) versus the conventional distinct infolding observed in BW (**Fig. 1V),** *cv.* Foma, and wild barley (*H. vulgare* subsp. *spontaneum*). This deviation was visible in all paleae independent of their position along the spike. Histological analyses using cross sections of paleae middle-areas (**Fig. 1U**) revealed distinct features of *com1.a* in which sclerenchymtous cells, in particular, appeared to be expanded in size and most likely also in numbers (**Fig. 1W-X**); however, we did not determine cell numbers quantitatively. Cell expansion is thought to be tightly linked to cell wall extensibility ^19,20^. We used transmission electron microscopy (TEM) to verify whether *com1.a* palea cells had altered cell wall features. Notably, mutant palea cells had clearly thinner cell wall structures, thus fewer mechanical obstructions for cell expansion, implicating that COM1 functions as a regulator of cell growth via cell wall modifications (**Fig. 1Y-Z** **and Supplementary Fig. 2**). Moreover, mutant paleae generally formed three vascular bundles (VB) (**Fig. 1W**) compared with two VBs in BW (**Fig. 1X**). By analogy to changes in palea cell walls, such alterations might also explain the rescission of SM identity, providing that COM1 similarly affects cell wall integrity in meristematic cells, e.g. SM cells or boundary cells (cells separating inflorescence meristem, IM, from SMs) (see below).

### *COM1* encodes a class II, subclass CYC/TB1 TCP transcription factor

To investigate the genetic basis of the *com1.a* phenotype, we constructed a genetic map by screening ~6,000 gametes for recombination events in an F_2_ population (Bowman × *com1.a*) followed by further analysis of F_3_ families. Thus, 15 critical recombinant F_2_-derived F_3_ families (i.e., 16 plants per family) were further analyzed (**Supplementary Table 1, 2 and 3; Supplementary Fig. 1C-F**). This delimited a ~1.4 Mb interval carrying eight genes, one of which is a predicted transcription factor (HORVU 5Hr1G061270) entirely absent in *com1.a*, likely due to an induced deletion (**Fig. 2A**). The remaining seven genes either were not expressed or not differentially regulated between Wt and *com1.a* mutant (see below, the transcriptome analysis). To validate our candidate gene, we sequenced it in a set of 20 induced barley spike-branching mutants and in a barley TILLING population of two-rowed barley cv. Barke. Resequencing of branching mutants, using both CDS and promoter specific primer pairs (**Supplementary Table 1**), revealed that five of them, i.e. *Mut.3906*, *int-h.42, int-h.43* and *int-h.44*, and *com1.j*, lost the same transcription factor as was found missing in the *com1.a* mutant (**Fig. 3**; **Supplementary Table 4**). All five mutants also showed the flat-palea phenotype observed in the mutant *com1.a* (**Fig. 3**). Allelism tests of *com1.a* with *Mut.3906* indicated that they are allelic to each other. Furthermore, PCR-screening of the TILLING populations for the CDS of the candidate gene revealed four homozygous M3 plants (M3.15104, M3.4406, M3.13729 and M3. 2598) carrying SNP mutations inside the DNA binding domain (**Fig. 2B**). Additionally, two heterozygous M3 lines M3.4063 and M3.9299 with SNP mutation outside the domain were also identified (**Fig. 2B**). All six SNP mutations caused amino acid substitution in conserved positions (**Fig. 2B**). They all transmitted a branched spike as was revealed by the phenotypes of the corresponding M4 and M5 homozygous plants (**Fig. 4**; **Supplementary Figs. 3 and 4; Supplementary Table 4**). Interestingly, from the six TILLING mutants, only two, with mutation within the TCP domain, showed either a true flat - palea phenotype with a complete loss of the infolding (line 2598, exhibiting also the most severe branching), or only a mild change in the palea shape (line 4406) (**Fig. 2B**; **Fig. 4**). Thus, penetrance of the mutant flat-palea phenotype depended on the type and position of the amino acid substitution (**Supplementary Fig. 3**, legend). Together, these data confirmed unambiguously that the transcriptional regulator was responsible for the spike-branching and palea phenotypes in *com1.a*. Annotation analysis of the COM1 protein showed that it belongs to the plant-specific TCP (Teosinte branched 1 (TB1)/Cycloidea/Proliferating Cell Factor) transcription factor family; COM1 contains 273 amino acids and features one basic helix-loop-helix TCP domain (**Fig. 2B**). Proteins of the TCP family fall into two classes, with COM1 belonging to class II, subclass CYC/TB1 ^21,22^.

**Fig. 2.**
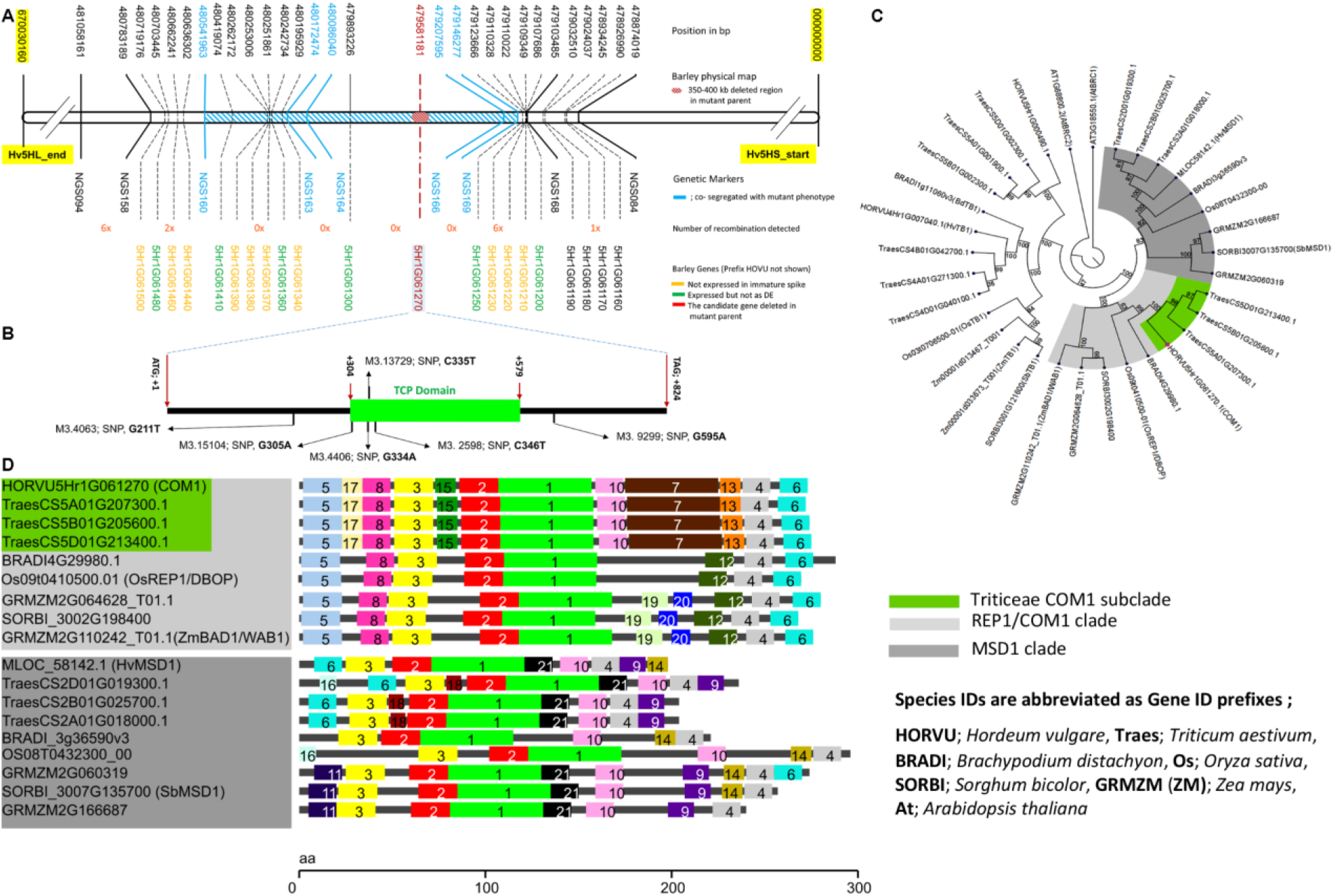
Map-based cloning of the gene underlying *com1.a*, phylogenetic analysis and protein structural variation of COM1. (**A**) Physical and genetic maps of *com1.a* from 100 recombina nt plants or ~6,000 gametes. A single gene (red; HORVU5Hr1G061270, a single-exon TCP transcription factor) was the strongest candidate and deleted in the mutant parent *com1.a*. (**B**) *COM1* gene model containing one TCP DNA binding domain (green box). Six barley TILLING alleles are shown with prefix M3. **(C)** UPGMA phylogenetic tree, using 1000 bootstrap replications, of COM1 homologs (highlighted in light gray) and paralogs (in dark gray) appeared as first- and second-best hits, respectively, in the blast search. Bootstrap values (in percentage) are shown within the circular cladogram along the edges of the branches. (**D**) Evolutionarily conserved motifs, among COM1 homologs and paralogous proteins (presented as phylogenetic tree in Fig. 2C), using the tool SALAD. Each colored box represents a different and numbered protein motif. For example, motif 1 in light green represents the TCP domain. Motifs 7, 13, 15 and 17 of the REP1/COM1 clade are specific to the Triticeae. (see also **Supplementary Fig. 6**).

**Fig. 3.**
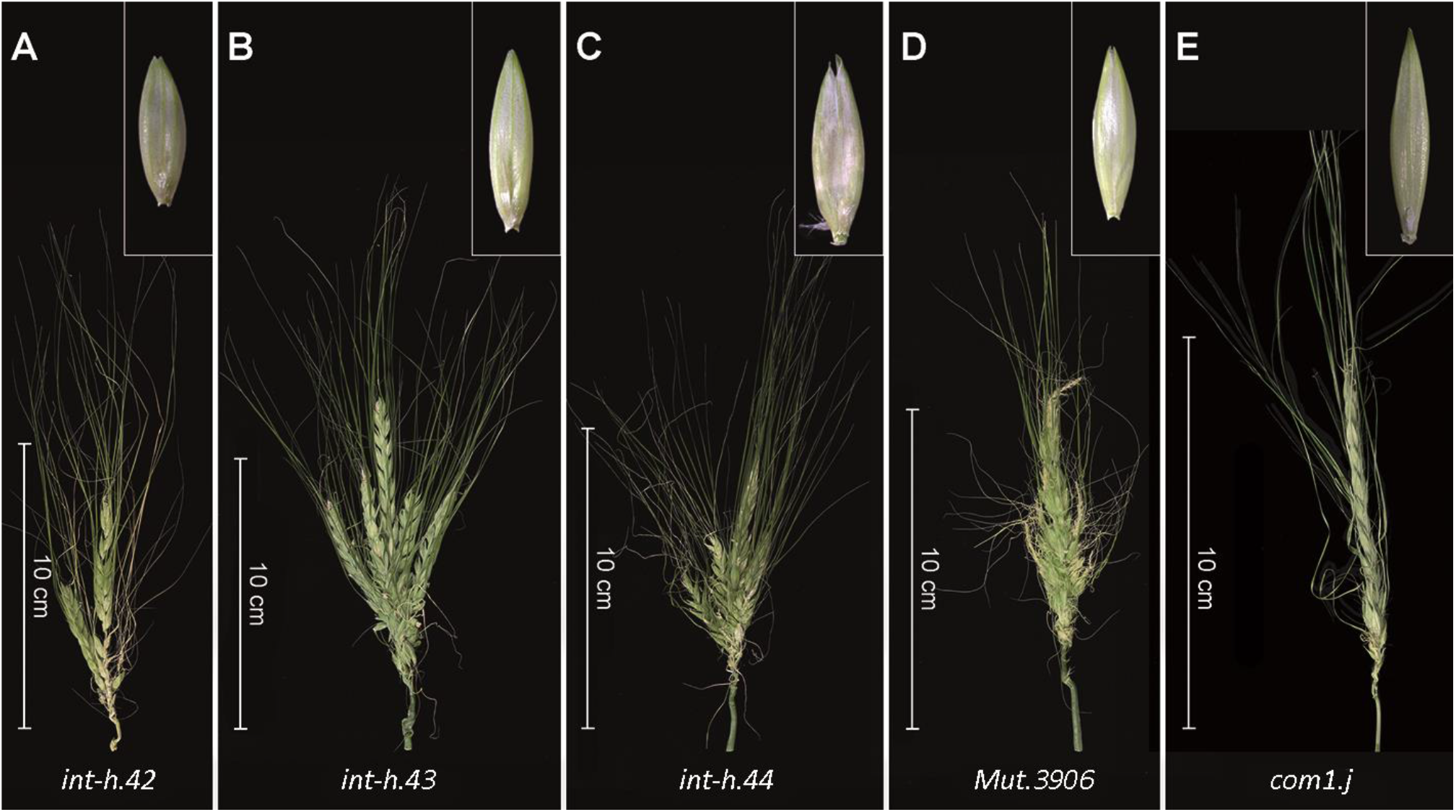
Barley *COM1* mutant alleles showing spike-branching and paleae phenotypes. Different induced mutant alleles identified by resequencing of primers correspond to CDS and putative promotor region of *COM1*. The corresponding palea is shown in the upper–right side of each spike image. See also **Supplementary Table** 1 and 4.

**Fig. 4.**
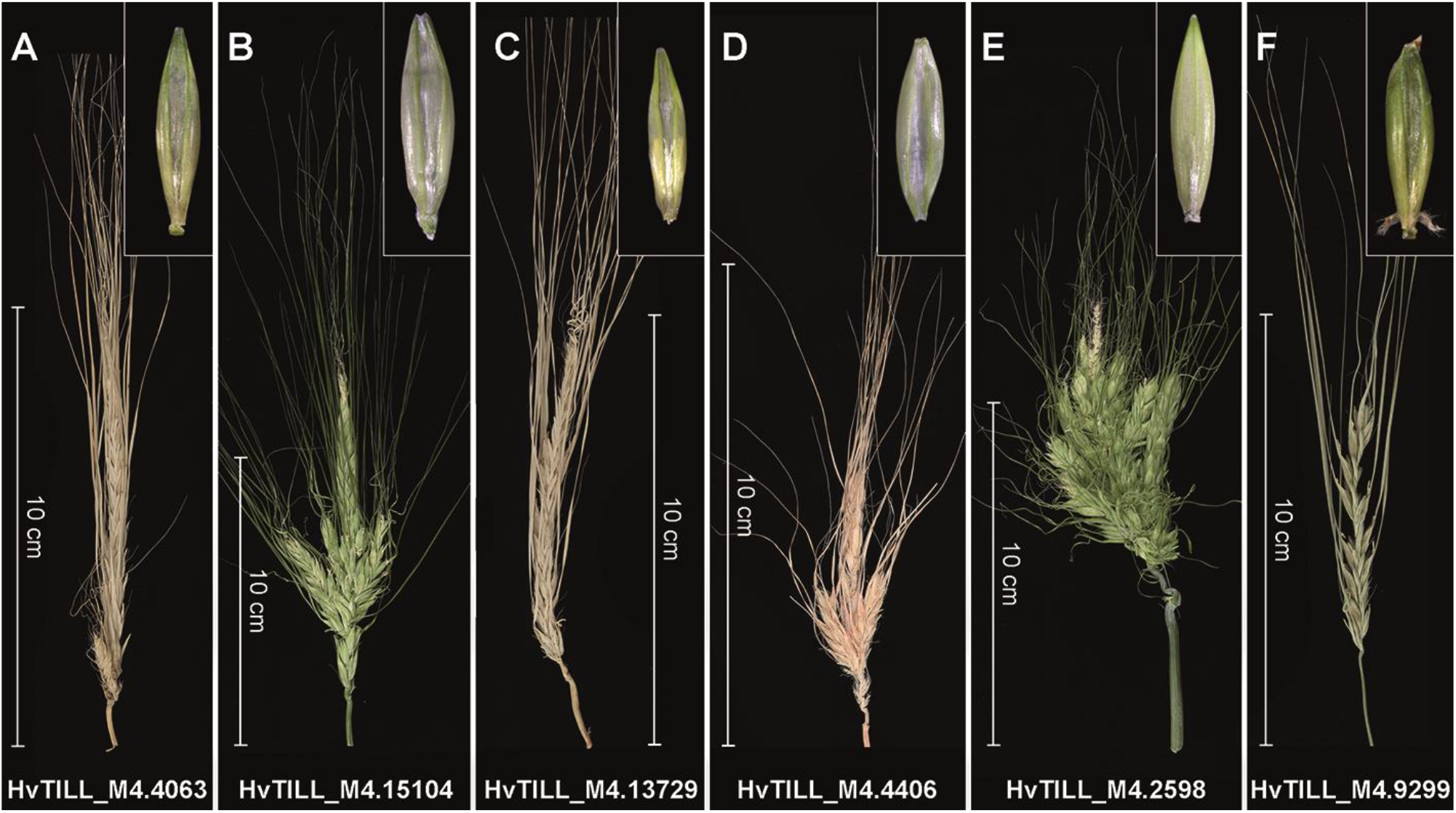
Spike and paleae phenotypes of barley TILLING lines. A representative display of branch formation of the six barley TILLING mutant plants derived from barley *cv.* Barke. The corresponding palea is shown in the upper -right side of each spike image. For the underling protein sequence lesion, see **Supplementary Figure 3**.

### Barley COM1 function evolved to affect boundary signaling

We next asked whether COM1 has experienced functional conservation or divergence within the grasses and whether its sequence composition supports possible functional alteration. We used the comprehensive phylogenetic analyses available for grass TCPs ^22,23^ (and the references therein) as a starting point for our own COM1-specific phylogenic analyses. We searched for homologs and paralogs of COM1 in sequenced grass genomes, including rice, maize, sorghum, hexaploid wheat and *Brachypodium distachyon*, as well as *Arabidopsis thaliana* (**Fig. 2C**). The homolog of maize TB1, obtained from the aforementioned grasses, was added as an out-group to the phylogeny. Our sequence searches and the phylogenetic analysis confirmed that COM1 is restricted to grasses (**Fig. 2C**) as reported previously ^14,15,24^. The homologs of COM1 in maize and rice were reported previously as *ZmBAD1/WAB1* and *OsREP1/DBOP* (60.3% and 65.5% sequence similarity to COM1), respectively ^14–17^. Except for maize, none of the COM1 homologs showed a duplication after speciation (e.g. no in-paralogs resulting from within-genome duplication, ^25^). Furthermore, COM1 seems to be a paralog (e.g. out-paralog ^25^ that refers to duplication before speciation; see **Supplementary Fig. 5A-B**) of the sorghum gene *SbMSD1* (44.1% sequence similarity to COM1) ^*26*^. Functional characterization of COM1 homologs is only available for maize and rice (Table 1) ^14,15,17^.

**Table1.** Functional variation of COM1 homologs observed among grass species.

| Species (Name) | Gene function in boundary... | Effect on the corresponding organ/meristem |  |  |  |  |
| --- | --- | --- | --- | --- | --- | --- |
|  |  | On branch formation | On pulvinus size/formation | Growth of palea cells | Number of VB<sup>1</sup> in palea | Pollen fertility<sup>2</sup> |
| Barley (COM1) | signaling | Inhibition | absent<sup>3</sup> | Inhibition<sup>4</sup> | Promotion | Normal |
| Brachypodium (BdWAB1/BAD1) | formation<sup>5</sup> | No effect | Promotion | No effect<sup>6</sup> | Promotion | Normal |
| Rice (REP1) | formation | Not reported<sup>7</sup> | Promotion | Promotion | Promotion | Reduced |
| Sorghum (SbWAB1/BAD1) | formation | Promotion | Promotion | No effect | Promotion | Reduced |
| Maize (WAB1/BAD1) | formation | Promotion | Promotion | Not reported | Not reported | Not reported |

<sup>1</sup> stands for Vasculature Bundles
<sup>2</sup> revealed by grain setting measurements as a proxy
<sup>3</sup> pulvinus is typically absent in Wt spike of Triticeae including barley as well as in the branched mutant spikes
<sup>4</sup> apparent at the longitudinally-middle palea part resulting in the formation of the infolding
<sup>5</sup> Refers to the formation of a boundary between pulvinus and the lateral branch without which fusion of the two happened; reflects intermediate evolutionary phylogenetic position of Brachypodium among grasses
<sup>6</sup> not visible at the microscopic level
<sup>7</sup> perhaps because the rice cultivars used in the corresponding studies (*cv. Nipponbare* and *cv. 9522*) are known to exhibit panicles with acute lateral branches.

Maize *BAD1/WAB1* transcripts are mainly detected at the IM-to-BM (branch meristem) boundary region as well as between pulvinus and lateral branches (in Fig. 3J of ^15^). Consequently, loss-of-function *bad1/wab1* mutants display organ fusion (a known boundary formation defect) resulting in reduced branch number (from 5.8 in Wt to 1.3 in mutant siblings) and angle size, and more upright tassel branches ^14,15^. This gene was dubbed a boundary formation gene promoting lateral meristem (e.g. branch) and axillary organ (e.g. pulvinus) formation in Wt maize ^14,15^.

Our phylogenetic analysis identified orthologs of *COM1*, in both sorghum and *Brachypodium distachyon* (**Fig. 2C**). Thus, to further expand our knowledge about *COM1* function within non-Triticeae, we therefore studied these species using a TILLING approach. We first screened a TILLING population in sorghum originated from *cv.* BTx623. The sorghum Wt inflorescence, a panicle, consists of a main rachis on which many primary, secondary as well as sometimes tertiary branches develop (**Fig. 5A**). Similar to maize, sorghum plants possess a pulvinus to regulate branch angle. The TILLING analysis revealed one mutant (ARS180 line; A144T) with both upright panicle branches (10.95° in Wt vs. 5.2° in mutant, *P*≤0.001; **Fig. 5A-G**; **Supplementary Table 5**) and reduced primary branch number per “node” e.g. whorls of branches (5.1 in Wt vs. 4.3 in mutant, P≤0.05; **Supplementary Table 5**). Measurement of branch angle was used as a proxy for pulvinus development (**Supplementary Table 5)**. These data suggest a similar positive role of sorghum *BAD1/WAB1* in pulvinus development and branch initiation/formation, revealing functional conservation of the protein between sorghum and maize. Moreover, we detected no obvious change in sorghum palea morphology except one additional vascular bundle, similar to maize and barley (**Table 1**).

**Fig. 5.**
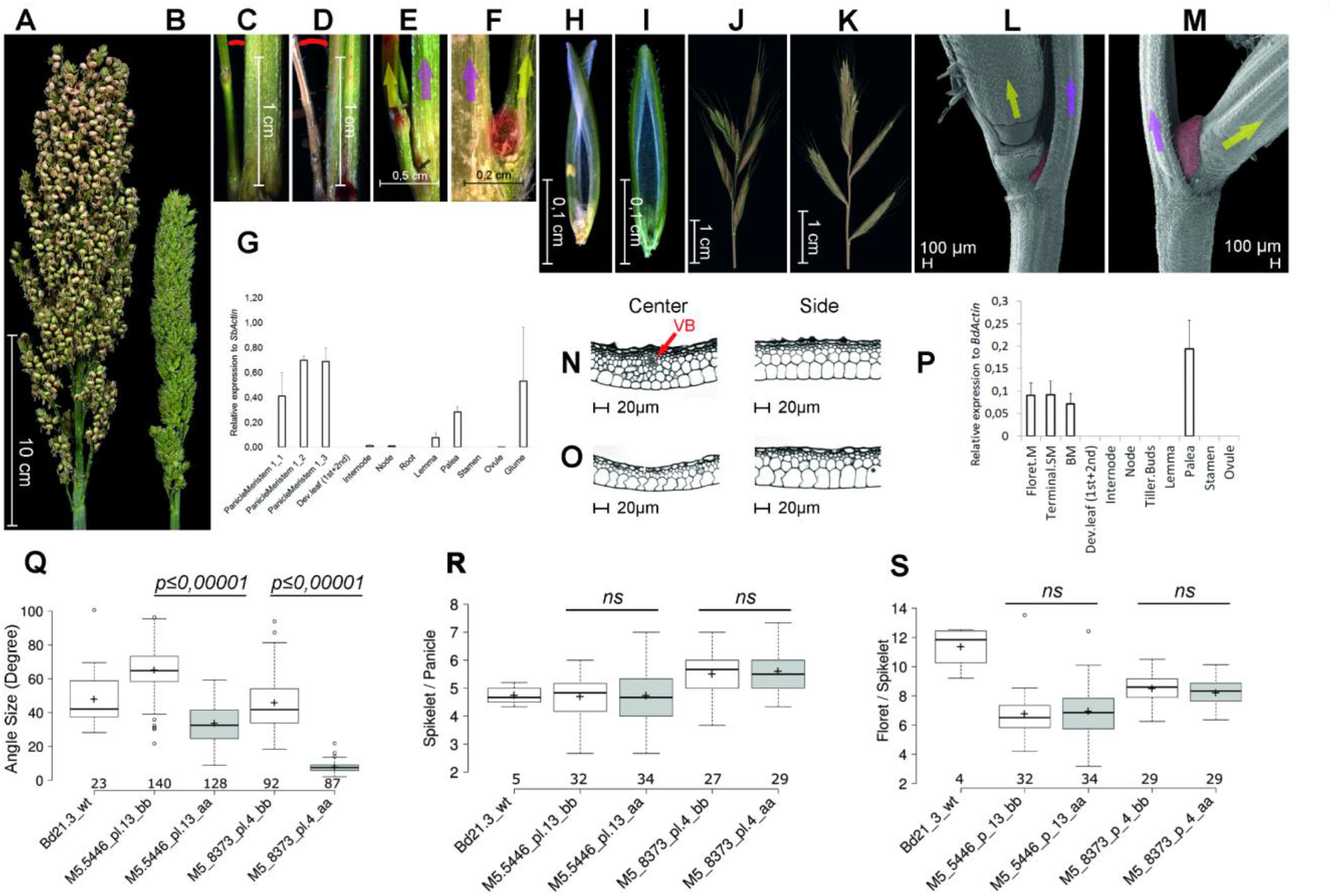
Inflorescence morphology and gene expression patterns in sorghum (A-G) and Brachypodium (H-S). (**A**) Sorghum inflorescence shape in Wt *cv.* BT623. (**B**) Compact sorghum inflorescence in TILLING mutant ARS180 showing severe reduction in grain setting (Supplementary Table 5) as also reported for *rep1* mutant in rice (see Table 1). (**C**) Acute branch angle in mutant ARS180 versus (**D**) Expanded branch angle of Wt (5.2° in mutant vs. 10.95° in Wt, *P*≤0.001; see Supplementary Table 5). due to the lack or small size of the pulvinus. (**E**) Depicts lack of pulvinus at the base (black arrow) of the mutant lateral branch versus its presence (red, roundish area) in Wt (**F**). Arrows in yellow and pink represent the lateral primary branch and rachis, respectively, in both E and F. (**G**) RT-qPCR of *SbBad1/Wab1* in organs of Wt plants. 1_1, 1_2 and 1_3 represent first, second and third branch meristem stages, respectively. **(H)** Brachypodium mutant palea scissor-like structure collapses easily due to external mechanical pressure; **(I)** normal/solid palea structure in Wt plants. **(J)** Brachypodium mutant inflorescence with compact shape due to acute branch (spikelet) angles; **(K)** Brachypodium Wt inflorescence with normal architecture of expanded branch angle as result of normal growth with pulvinus. (**L**) SEM view of an abnormal tiny pulvinus (in red) of a Brachypodium mutant versus an intact normal-sized pulvinus (in red) in Wt (**M**); arrows in yellow and pink represent the lateral branch and rachis, respectively, both in L and M. (**N**) Histological view of transverse section of Brachypodium mutant palea as compared to Wt (**O**); Brachypodium mutant has an extra VB in the center part (red arrow) which is lacking in Wt. Center refers to the collapsed middle part while Side refers to the flanking intact area (the blades of the scissor; see part H). (**P**) RT-qPCR of *BdBad1/Wab1* gene expression across meristematic stages and organs in Wt. **(Q)** Branch angle measurement in Brachypodium as proxy for pulvinus size. **(R)** Number of spikelets per individual Brachypodium infloresce nce (panicle). **(S**) Number of florets per spikelet in Brachypodium. In Q to S; data are from contrasting M6 homozygous TILLING lines of Brachypodium; aa and bb refer to homozygous mutant (aa) and Wt (bb)a alleles from the same family (**Supplementary Note**). Values above x-axis indicate number of items used to collect data points (representing number of angles measured in Q, and number of plants in R and S). *P* values were determined by using *Student’s t* test; ns: not significa nt. Genotype IDs below x-axis refer to the parental line of the respective M6 family. For Q-S; Source data are provided as a Source Data file.

The rice homolog of *COM1*, *OsREP1/DBOP*, shows a major effect in promoting palea identity, growth and development, with no effect on branch angle or branch initiation ^16,17^. Loss-of-function mutants display smaller paleae due to less differentiation and severely reduced size of palea cells; a clear contrast to palea defects in barley (**Table 1**). Our TILLING analysis of COM1 homologs in *Brachypodium distachyon* (for its Wt inflorescence shape see schematic **Fig. 1C** and **Fig. 5K**) identified several mutants. Phenotypic investigation of two lines (5446: Q116* and 8373: S146N) (**Supplementary Table 4, Supplementary Note**) revealed similar phenotypes to the aforementioned non-Triticeae species (**Table 1**) (**Fig. 5H-P**). Similarly, we observed a palea defect (**Fig. 5H-I**) but histological analyses revealed no changes in cell expansion, except the formation of one additional vascular bundle in each mutant (**Fig. 5N-O**). We also observed a reduction in branch angle because of smaller or absent pulvini (**Fig. 5J-M**); however, the number of lateral branches was not altered in either Brachypodium mutants (**Fig. 5Q-S**). In conclusion, COM1 homologs within non-Triticeae grasses primarily promote boundary formation and cell differentiation (as in rice palea)/proliferation (as seen for pulvinus) (**Table 1**); but similarly promote the formation of lateral axillary organs, e.g. branch or pulvinus, to contribute in maintaining complex inflorescence structures.

To better understand the contrasting COM1 function of branch-inhibition in barley versus branch-formation in non- Triticeae grasses, we analyzed barley *COM1* expression using qRT-PCR and semi-qPCR (Fig. 6A-C) followed by mRNA *in-situ* hybridization (**Fig. 6D-G**). Barley *COM1* transcripts were detected in paleae (**Fig. 6C, F-G**), VB of the rachis (Fig. 6E), and importantly at the base of forming SMs throughout the boundary region separating SMs from IM (IM-to-SM boundary) and between lateral and central SMs (**Fig. 6E-F**), similar to non-Triticeae grass species, e.g. maize. This expression pattern suggests involvement of barley *COM1* in specification of the spikelet meristematic boundary. However, since central and lateral spikelets do not fuse into each other or to the IM (as long branches do fuse to the IM in maize or sorghum), barley COM1 may not be involved in boundary formation *per se* but perhaps rather in boundary signaling (see below transcriptional result and discussion). ^27^. Recently acquired protein motifs specific to Triticeae COM1 may support this functional difference (**Fig. 2D** Motifs 7, 13, 15 and 17 **and Supplementary Fig. 6**).

**Fig. 6.**
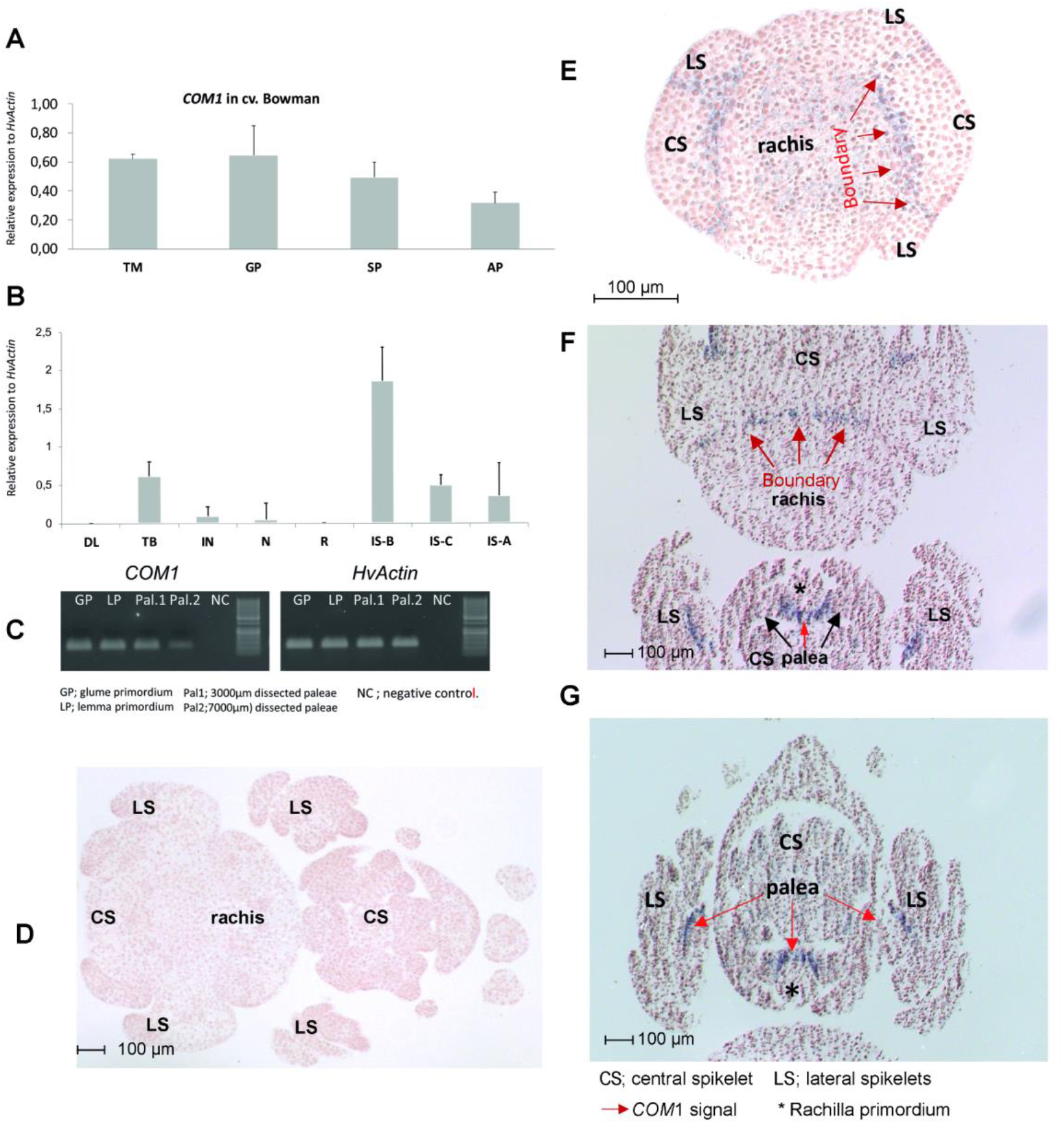
Transcript analyses of *COM1* in two-rowed barley. **(A)** Relative *COM1* expression at different stages of immature barley spike including TM, GP, SP as well as the at meristematic stage of awn primordium (AP; a stage following stamen primordium ^18^) in cv. Bowman. (**B)** Relative *COM1* expression in different organs (DL; developing leaf, TB; tiller buds, IN; culm internode, N; culm node, R; Root) along with spike sections (IS-B; immature spike basal nodes, IS-C; immat ure spike central nodes, IS-A; immature spike apical nodes) at meristematic stage of AP in *cv.* Bowman. Despite expression in tiller buds, no differences in tiller number was observed (**Supplementary Fig. 4**). Dev leaf and IM stands for developing leaf and inflorescence meristem, respectively. (**C)** Semi-qPCR of *COM1* (left) and *HvActin* (right) mRNAs in two different stages of immature spike development, GP and LP (as positive controls) as well as in two palea samples. **(D)** *COM1* mRNA in-situ control hybridization using pooled sense probes (see online methods). (**E-G**) mRNA in-situ hybridization of *COM1* using pooled anti-sense probes. Tissues represent cross-section through a spikelet triplet at TM **(E)** and AP stages (**F-G**) of barley *cv.* Bonus (a two-rowed Wt). For D-G; Source data is provided as a Source Data file.

We checked whether natural selection has acted upon barley COM1 sequence composition and function, and consequently formation of unbranched spikes in barley. Re-sequencing of the barley *COM1* coding sequence in a panel of 146 diverse barley landraces and 90 wild barleys ^28,29^ revealed very little natural sequence variation (site diversity of pi = 0.0006). Eleven SNPs resulted in a simple 12-haplotype network (**Supplementary Fig. 7**) comprising only two main haplotypes, neither of the 12 showed mutant spike or palea phenotypes (**Supplementary Fig. 7**). This suggests that barley COM1 underwent purifying natural selection. We assume that this selection may have contributed to maintaining barley’s slimmed-down inflorescence shape.

### *COM1* inhibits inflorescence branching partially independent to *COM2*

We had previously reported a branch suppressor gene, the AP2/ERF transcription factor *COM2*, with a conserved function across grass species ^13^. *COM2* expresses in an arc-like region between central (CS) and lateral spikelets (LS) as well as between the SM and the emerging GP ^13^. As the *com1* phenotype resembles that of *com2 (***Fig. 7A-C**), we developed and characterized BW-NIL(*com1./com2.g*) double mutants (e.g. DM, or *com1./com2.g* double mutant) to study their interactions in regulating branch inhibition in barley.

**Fig. 7.**
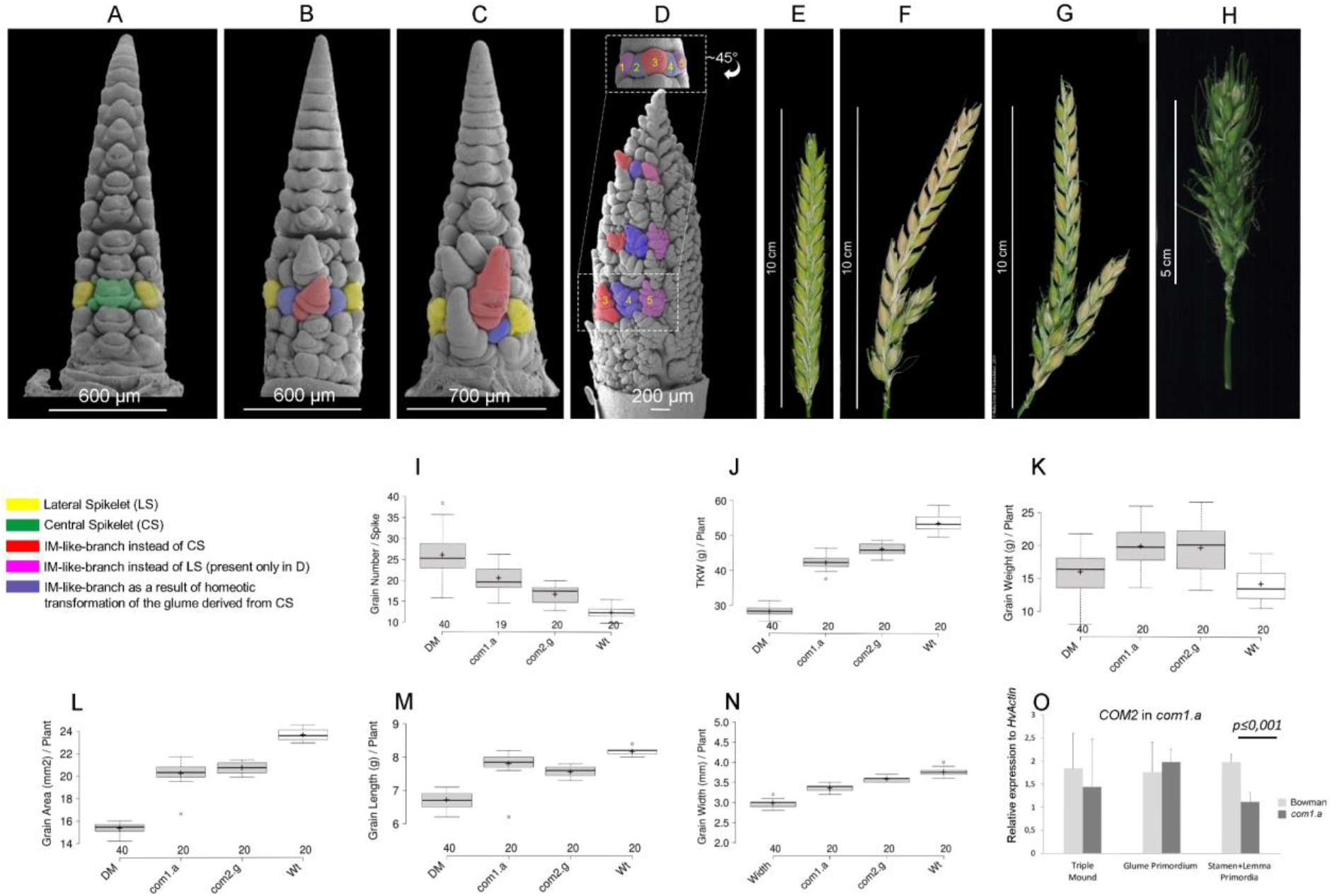
Immature and mature barley spike morphology in wild type versus single and double mutants (A-H), with their comparative grain-related characters (I-N) as well as the transcript levels of *COM2* in single mutant *com1.a* **(O).** (**A**) SEM-based view of an immature spike of Wt *cv.* Bowman, single mutant *com1.a* (**B**), single mutant *com2.g* (**C**) and those of a DM mutant (**D**). In **D,** upper panel shows basal nodes of a DM spike at GP stage with elongated CSM (as compared to Wt in Fig. 1G) and unusually enlarged glume primordia (in purple). Numbers 1 to 5 denote five IM-like branch meristems of a DM spike that eventually represents a putative ten-rowed spike. The lower panel is rotated by ~45° to the left while imaging. Please note; IM-like branches 1 and 2 are not visible in the lower panel. Immature spikes (except D upper panel) are at similar developmental stages of early/advanced stamen primordium. Immature spikes in B and C represent a typical SBS while D corresponds to the immature CBS spike class. (**E-H**) Depicts mature spikes in Wt *cv.* Bowman (**E**), single mutant *com1.a* (**F**), single mutant *com2.g* (**G**) and the mature spike in a DM plant (**H**). **F** and **G** represent a frequent phenotype of the SBS class at maturity while **D**represents a mature CBS phenotype. (**I-N**) Grain characters of the DM plants, and the corresponding single mutant *com1.a* and *com2.g* in comparison to the Wt *cv*. Bowman. Data are based on a single greenhouse experiment and on averages of 20 plants (390 to 540 spikes) per phenotypic class. (**O**) Depicts *COM2* transcripts in the *com1.a* mutant compared to Wt *cv.* Bowman. Mean ± SE of three biological replicates per stage are shown. Genotype differences were tested at a significance level of P > 0.05. For I-N; Source data are provided as a Source Data file.

We first performed a comparative SEM-based image analysis in immature spikes of the two singles and the DM mutants. As illustrated above, SEM of the two single mutants revealed similar branched phenotypes by generating a simple branch structure (SBS) in which the CSMs located at the more basal nodes lost identity and converted into IM-like branches (for a typical SBS see**; Fig. 7B-C)**. SBS also included homeotic transformations of glumes **(Fig. 7B-C**, in purple). In contrast, DM immature spikes revealed interesting observations including loss of identity and conversion of LSMs to IM-like branches, in addition to the loss of identity in CSMs **(Fig. 7D)**. These conversions were observed in all nodes; not only in basal ones. Furthermore, glume primordia that underwent only an occasional homeotic transformation in basal nodes in single mutants (in purple, **Fig. 7B-C**), were also converted into IM-like meristems at all nodes of a DM spike (in purple, **Fig. 7D**). Therefore, a mixed meristematic re-organization per immature DM spike was observed representing a tentative ten-rowed barley spike (**Fig. 7D** and the legend), and thus, a rather complex branch structure (CBS).

To further characterize the three genotypic classes, i.e. DMs, *com1.a* and *com2.g,* we also compared spike morphology at maturity. A set of 20 plants per class as well as 20 wild type *cv.* Bowman plants were grown to perform a comparative phenotyping of mature spikes. Our visual inspection of the two single mutants at maturity revealed three types of spike forms including Wt (**Fig. 7E**), SBS (**Fig. 7F-G**; typical SBS branching forms) and CBS (**Fig. 7H**). In case of Wt inflorescence architecture, *com1.a* displayed only 3,7% of the spikes per family in this class, while *com2.g* showed a higher frequency of 22%. As expected, SBS was the most frequent class in both single mutant families with 91% and 73% of spikes per family in *com1.a* and *com2.g,* respectively. Interestingly, both single mutants showed also a low level of CBS with similar frequency (*com1.a;* 5,3%, *com2.g;* 5,1%) that was mostly visible in small late tillers. Thus, *com1.a* mutant showed a higher phenotypic penetrance for spike-branching, e.g. higher level of SBS and lower frequency of Wt spikes, as compared to *com2.g* mutant plants. In contrast to the single mutants, all (100%) spikes of the DMs displayed the CBS class (**Fig. 7H**), leading DM plants to outperform either single mutant in supernumerary spikelet formation, and thus, in grain number per spike (**Fig. 7I)**. We further measured other grain-related characters (**Fig. 7J-N**), showing that the DMs had the lowest TKW; most likely due to the known trade-off with increased grain number.

To further examine the genetic interactions between *COM1* and *COM2* during branch inhibition of the barley spike, we performed qRT-PCR analyses. *COM2* transcript levels in immature spikes of *com1.a* were unchanged during the two early stages tested; however, slightly lower expression was only found during later stages of development (**Fig. 7O** **and** **Fig8A**; dashed red arrow). Thus, the DM analyses imply that the two loci may act partially independently/additively during branch inhibition in barley.

**Fig. 8.**
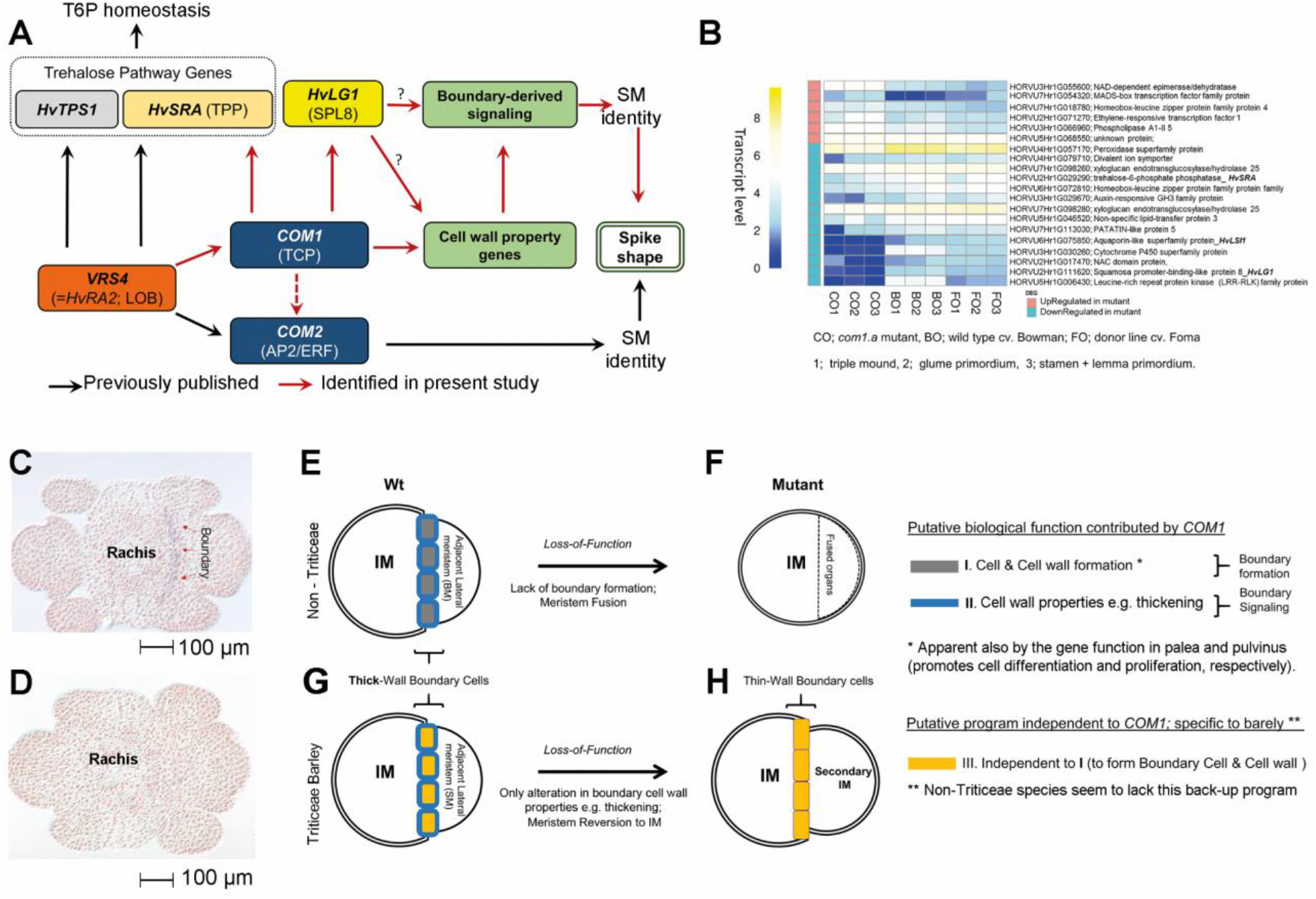
Model of *COM1* regulation based on transcriptome analysis in barley (A-B), transcript analyses of *HvLG1* in two-rowed barley (C-D), and schematic representation of functional *COM1* differences from non-Triticeae (E-H). Model of *COM1* transcriptional regulation deduced from either RNAseq or RT-qPCR results. Black arrows are interactions reported previously ^13,28^ while red arrows are detected in the current study (see Supplementary Fig. 10, the legend). **(B)** RNA-seq-based heat map of selected differentially expressed (DE) genes (see **Supplementary Fig. 9** for the remaining DE genes). The transcript level of each gene in mutant *com1.a* (CO) is compared with two Wts, cv. Bowman (BO; parent of the mapping population) and cv. Foma (FO; the donor line), at three different meristematic states. Transcript level= Log(X+1)-Scaled Expression; where X is the normalized expression value of a given gene (see online Methods). **(C)** mRNA in-situ hybridization of barley *LG1* in the immature spike using the antisense **(D)** and sense probes (see online methods). Tissues represent cross-section through spikelet meristems in barley *cv.* Bonus at GP stages. **(E-H)** Proposed IM-to-BM boundary formation due to Wt gene function in non-Triticeae grasses (**E**). Lack of boundary formation due to the loss-of-function allele (**F**). Of note, involvement in the alteration of the boundary cell walls within *non-* Triticeae species cannot be excluded. (**G**) Proposed IM-to-SM boundary formation in Wt barley; restriction of COM1 function to altering cell wall properties (the blue program), due to evolutio nary functional differences. (**H**) Reversion to the previous identity state (IM) observed in barley *com1.a* due to lack of putative wall-amplified micromechanical signals needed to confer SM identity. For A-B as well as C-D; Source data are provided as a Source Data file.

### Putative transcriptional regulation during barley spike development

To further examine the molecular basis of COM1 branch inhibition within the barley spike, we performed qRT-PCR to locate *COM1* relative to other previously known spike architecture genes (**Fig. 8A**, black arrows). In addition to the interactions with *COM2* (Fig. 7O), we localized *COM1* downstream of *VRS4 (HvRA2*; orthologous to maize *RAMOSA2*), the main regulator of row type and branch inhibition (**Supplementary Fig. 8A-D)** ^7,13^using qRT-PCR analyses of *COM1* expression in the BW-NIL(*vrs4.k*) mutant (**Supplementary Fig. 8E and** **Fig. 8A**).

We performed comparative RNA-seq using mRNAs from immature spikes of BW and *com1.a* as well as the mutant progenitor, *cv.* Foma, when spike patterning begins to differ between genotypes, plus two subsequent stages (**Figs. 1** **and** **8B**; **Online Materials**). Differentially expressed (DE) genes were identified in comparisons of *com1.a* versus BW and mutant versus *cv.* Foma. We found 83 genes (Log2 FoldChanges; LFC | ≥ 0.5; adjusted *P* < 0.05) DE in at least one stage in both comparisons (**Fig. 8B**; **Supplementary Figs. 9–10; Supplementary Source Data**): 18 and 65 genes up- and downregulated in BW-NIL(*com1.a*), respectively.

Among significantly downregulated genes across all three stages (**Fig. 8B**), we detected one *SQUAMOSA PROMOTER-BINDING-LIKE 8* gene (*SPL8*, HORVU2Hr1G111620) homologous to the boundary gene *LIGULELESS 1* in maize (*LG1*; *ZmSPL4*), rice *OsLG1* (*OsSPL8*) and hexaploid wheat *TaLG1* (*TaSPL8*) ^30^. Similar to the known maize module (*RA2*→*WAB1*/*BAD1*→*LG1*; ^14,15^, we found that *VRS4*/*HvRA2*→*COM1*→*HvLG1* regulation appears to be maintained in barley. *HvLG1* mRNA *in-situ* hybridization showed co-localization with *COM1* in the base of the forming SMs throughout the boundary region separating SMs from IM (IM-to-SM boundary) (**Fig. 8C-D**). Transcriptome analysis of leaf tissues in a wheat *liguleless1* mutant revealed *TaSPL8* as a cell wall-related gene ^30^. Notably, no spike-branching phenotype was reported for this erected-leaf *liguleless* mutant, most likely due to genetic redundancy.

Among other significantly downregulated genes in *com1.a*, we found important genes associated with cell wall properties and integrity (**Fig. 8B**; **Supplementary Fig. 9**). These include HORVU5Hr1G006430, a leucine-rich repeat receptor kinase (LRR-RLK), and HORVU3Hr1G030260 belonging to the cytochrome P450 superfamily. LRR-RLKs and CYP450s are involved in lignin deposition to cell walls upon cellulose biosynthesis inhibition and during lignin biosynthesis *per se*, respectively ^31,32^. Other cell wall-related genes include two genes encoding xyloglucan endotransglucosylase/hydrolase (XTH) 25 (HORVU7Hr1G098280 and HORVU7Hr1G098260) and barley *Low Silicon Influx 1* (*HvLSI1*; *HORVU6Hr1G075850*) ^33^, both downregulated in the mutant. These cell wall-related genes may support *COM1* involvement in regulation of cell wall mechanics of palea cells and the IM-to-SM boundary, and indirectly, putative signaling required for acquiring SM identity.

## Discussion

Here we report that barley COM1 affects cell growth through regulation of cell wall properties specifically in palea and IM-to-SM boundary cells; the latter provide identity signals to barley SMs ^34^. Signaling to the SM to establish its identity is a key genetic switch by which barley inflorescences acquire spike architecture, not seen in non-Triticeae grasses.

*COM1* is present only in grasses, with no true *Arabidopsis* ortholog; intriguingly, we observed functional differences of COM1 between barley and non-Triticeae grass species. The differences in COM1 function was clear by comparing mutant versus wild type inflorescence phenotypes across grass species, and was further elucidated by our analysis at the cellular/molecular level. At the phenotypic level, barley COM1 inhibits spike-branching to simplify floral architecture; whereas in non-Triticeae COM1 homologs promote formation of lateral branches (e.g. up to 60% more branches in maize when compared to mutants ^15^) to sustain the ancestral inflorescence complexity.

At the cellular level in non-Triticeae grasses, COM1 has evolved as a boundary formation factor, its putative ancestral role (**Fig. 8E-F**). Consequently, loss-of-function of COM1 homologs result in lack of boundaries and subsequent organ fusion, e.g. BM into IM, as demonstrated by a low number of lateral branches in maize mutants. Notably, this loss of function did not change the overall inflorescence architecture in non-Triticeae grasses. Barley COM1 loss-of-function, however, increases branch formation/extension mostly from SMs, a clear deviation from the canonical spike form. As barley *COM1* displayed a similar boundary mRNA expression as seen in maize, we presume that barley COM1 functions through boundary signaling ^34^, thereby affecting the identity of adjacent SMs. The formation of boundary regions in barley *com1* mutants (no organ fusion) via pathway(s) independent of *COM1* (**Fig. 8G-H**), and thus separation of meristematic zones in this mutant, implies that barley IM-to-SM boundary cells fail to deliver proper identity-defining signals to SMs. This signaling failure may perturb transcriptional programs required to establish identity in barley SMs; such meristems eventually revert back to IM-like meristems forming a branch-like structure (**Fig. 5H)**. The function of the boundary, and boundary-expressed genes (e.g., maize *RAMOSA1–3*), as a signaling center for adjacent meristems, e.g. SMs, has been proposed in grasses, yet features of these signals remain unknown ^34^. Signals associated with COM1 might include micromechanical forces derived from formation of rigid cell walls enclosing boundary cells. Involvement of COM1 in printing such mechanical regulation is supported by our anatomical analysis of palea cell walls and further confirmed by our transcriptome analysis of immature barley spike samples. *HvLG1*, *HvLSI* and genes encoding one LRR-RLK, one CYP450 and two XTHs were among the most downregulated in the mutant and involved in defining cell wall properties ^30–32,35^. The contribution of boundary cell wall mechanics in guiding organogenesis within reproductive tissues has been well described in eudicot species^36,37^.

Such functional differences usually include constraints on expression patterns, protein sequence/structure or participation in molecular networks, often assumed to be associated with gene duplication ^25^. Notably, *COM1* shows no sign of duplication within the barley genome and as mentioned above displays a similar expression pattern to maize ^14,15^. Thus, COM1’s functional difference and implication in boundary-derived signaling seem to be associated with its protein sequence (**Fig. 2D**) and the respective downstream molecular networks. Furthermore, COM1’s role in regulating floral complexity-levels in grasses fits well with the view that TCP transcription factors are growth regulators and evolutionary architects of plant forms that create diversity ^38^. They influence the final architecture of plants in response to endogenous and/or external conditions. Thus, the barley floral reductionism (from compound spike to spike form; **Fig. 1A-D**) contributed by COM1, might be a response to the ecological expansion of the Triticeae into more temperate climates ^3^.

In summary, our findings enabled identification of a barley SM identity pathway, *VRS4* (*HvRA2*) → *COM1* → *HvLG1*, which works partially independent of *COM2* and inhibits spike-branching via boundary-defined signals (**Fig. 8A** **and Supplementary Fig. 10**). Our model of branch-inhibition in barley spikes opens a new window into grass inflorescence evolution and molecular crop breeding, and the elevated grain number per spike in *com1.a/com2.g* double mutants supports this notion.

## Methods

### Barley Plant material

The Nordic Genetic Resource center, the National Small Grains Collection (US Department of Agriculture), and the IPK gene bank were inquired to access ‘Compositum-Barley’ mutants (Supplementary Table 4). Bowman NIL carrying *com1.a* allele ((i.e., BW-NIL(*com1.a*); syn. BW189 or CIho 11333)), its two-rowed progenitor Foma and Wt barley cv. Bowman were used for phenotypic descriptions, whole genome shotgun sequencing (WGS) (see below) as well as SEM analysis. Plant material used to generate mapping populations is reported in the corresponding section for genetic mapping. For haplotype analysis, a core collection including of 146 diverse barley landraces and 90 diverse wild barleys were sequenced ^28,29^ (**Source data file for Supplementary Fig. 7**).

### Plant phenotyping

#### Barley

For phenotyping the mapping population, BW-NIL(*com1.a*), Bowman and the corresponding segregating populations (F_2_ and F_3_) were grown side by side under greenhouse conditions at the IPK. For a plant to be assigned as a branched spike mutant, spike shape at all tillers was visually inspected for presence of at least one extra spikelet at any rachis node. Grain related characters such as weight, number, etc. were also measured at harvest for the two parental lines of the mapping population. In case of phenotyping of the barley TILLING population (see below and the **Supplementary Table 4**), other induced mutants (**Supplementary Table 4**) as well as the BW-NIL(*com1.a*) / BW-NIL(*com2.g*) double mutants (see below), visual phenotyping for variation in palea structure was also applied in addition to the aforementioned phenotyping approach used for spike-branching in F_2_ and F_3_ progenies. In case of TILLING, from the six mutants for which the spike-branching phenotype was observed at M4, only three (carrying mutation inside the protein domain; M4.15104, M4.4406, and M4. 2598) were subjected for further study at M5 generation. For which, one M4 plant was selected from which 16 M5 plants were grown and phenotyped.

#### Brachypodium distachyon

An already published TILLING population and the corresponding Wt accession Bd21-3 were used for phenotyping ^39^. That included measurement of branch angle, as proxy for pulvinus size, spikelet number per spike, floret number per spikelet and palea structure. Hence, per M4 plants, only homozygous M5 plants either with mutant genotype aa (3 to 4 plants) or wild type bb (3 to 4 plants) were selected. Per M5 plants, 10 M6 plants were grown under greenhouse conditions at the IPK and used for measurement. Thus, 30 to 40 plants per group and for each plant angles of basal spikelets in main tillers were considered for measurement. To this end, spikes were first imaged and then imported to the ImageJ tool (https://imagej.nih.gov/ij/index.html) for angle measurement. In case of original wild type Bd21-3, five plants were grown and measured. The same set of plants and the corresponding spike images were used to calculate number of spikelets per spike and number of florets per spikelet. In case of palea phenotyping: paleae were visually inspected across all spikes per plant. We detected plants with paleae being sensitive to exogenous finger-pressure, and thus such plants were scored as mutants. A gentle finger-pressure led the mutant paleae to crash from the middle longitude-line so that a scissors-like structure was formed (**Fig. 3G**). The crashing was not evident in Wt plants even with severe exogenous hand-pressure.

#### Sorghum

An already published TILLING population and the corresponding Wt accession BTx623 were used for phenotyping ^40^. To measure primary branch number and angle, 5 to 8 plants, either M5 or M6 generations, per family including a Wt sorghum family cv. BTx623 were grown under greenhouse conditions at the IPK. Average primary branch (p. branch) number per panicle, e.g. per plant, was calculated by counting all p. branches that originated per each rachis node for the first 10 nodes (Supplementary Table 5). The node refers to the rachis area where whorls of branches emerge. The average p. branch number per family was then used to compare with the same value obtained from Wt family BTx623. To measure the branch angle, for each plant 3 to 4 basal nodes per panicle were separately photographed. Each node contained at least 1 and up to 5 lateral branches. To cover angles of each individual branch per node, each node was photographed multiple time. Images were then imported to ImageJ for angle measurement as described for Brachypodium (see above). Spikelet organs of palea and glume as well as overall grain set were also visually inspected for any visible alteration.

### Marker development

BW-NIL(*com1.a*) and two-rowed progenitor of *com1.a,* cv. Foma, were survey sequenced using WGS approach (see below). These sequence information were compared against already available WGS of Bowmann ^41^, as present in **Supplementary Fig. 1**. Polymorphisms e.g. SNPs detected from this comparison (named as Next Generation Sequencing based markers (NGS-based markers)) between the two parental lines were converted to restriction enzyme based CAPS (http://nc2.neb.com/NEBcutter2/) markers to derive a restriction based genetic marker as previously described ^13^. The developed genetic markers (**Supplementary Table 1**) were used to screen the corresponding mapping population.

### Genetic mapping and map-based cloning of *com1.a*

***com1.a*** was initially proposed to be located in chromosome 5HL with unknown genetic position ^12^. A barley F_2_ mapping population was developed by crossing Bowman introgression line, BW-NIL(*com1.a*), and barley cv. Bowman. For initial mapping 180 individuals were analyzed and genotyped using the aforementioned NGS based markers. The pattern of segregation between mutant and Wt F_2_ plants fitted a 3:1 ratio typical for a monogenic recessive gene. Linkage analysis of segregation data was carried out using maximum likelihood algorithm of Joinmap 4.0. Kosambi mapping function was used to convert recombination fractions into map distances. The linkage mapping was further followed by a high-resolution genetic mapping in which almost 6,000 gametes were screened with the flanking markers NGS045 and NGS049. For narrowing down the *com1.a* genetic interval; the identified recombinants (a set of 109) were used. From 109, a set 15 F_2_ were labeled (**Supplementary Table 2-3**) to be critical recombinants for precisely defining the *com1.a* genetic interval. From each of the 15 critical plants, 16 F_3_ progenies were evaluated for their phenotypes and marker genotypes at the *com1.a* candidate gene. (**Supplementary Table 2 and S3**). Based on F_2_ high-resolution mapping and F_3_ genetic analysis described, two tightly linked markers, NGS084 and NGS094, were taken to harvest the available barley genome BAC sequence data (data not shown). A single BAC contig spanning 1.4 Mb of the minimal tiling path (MTP) was identified. Genes in this region were utilized for marker development and further genetic mapping that resulted in identification of a ~380 kb region deleted in the mutant BW-NIL(*com1.a*). The deleted fragment contains a single gene, i.e., *com1.a.*

### Allelism test of *com1* mutants

Mut.3906 mutant (**Supplementary Table 4**) was crossed with BW-NIL(*com1.a*) to test for allelism. The resultant F_1_ plants showed a mutant spike phenotype confirming to be allelic with *com1*. All alleles showed phenotypic similarities with *com1* and mutations in the *COM1* gene sequences.

### Double-mutant analysis

Double mutants (DM) were generated by crossing mutant BW-NIL(*com1.a*) to BW-NIL(*com2.g*), followed by selfing of the F_1_ progeny. All obtained 183 F_2_ plants were subsequently genotyped (**Supplementary Table 1**). In case of *com2.g* mutation detection, a primer pair (Com2-Bw_SfiI_FR; **Supplementary Table 1**) spanning the A300C haplotype (that differentiate the Wt Bowman allele A from *com2*.g mutant C allele at position 300bp ^13^ were used for sequencing and to classify F_2_ genotypes for the *com2* locus. Thus, genotypic classes include C300C allele as homozygous mutant, AA as Wt and CA as heterozygous. In case of *com1.a*, a presence/absence marker was used (**Supplementary Table 1**), where absence of the *COM1* gene was considered as homozygous *com1.a* mutant. A total number of five plants were recovered as homozygous double mutants (from 183 F_2_ plants) that were used for generating F_3_ plants used in subsequent DM phenotypic analysis (**Fig. 7**). Two DM F_3_ families, each consisting of 20 plants along with 20 plants from each of the single mutants and 20 wild type cv. Bowman plants, were grown and used for phenotyping (**Fig. 7**).

### TILLING analysis

#### Barley

For identifying further mutant alleles of *COM1* in barley TILLING populations including EMS (Ethyl methanesulfonate) treated population of cv. Barke consisting 10279 individuals were screened ^42^. A primer combination (**Supplementary Table 1**) was used to amplify the coding region of the *COM1* gene. The amplicon was subjected to standard procedures using the AdvanCETM TILLING kit as described in ^13^. Amplified products were digested with dsDNA cleavage kit followed by analysis via mutation discovery kit and gel-dsDNA reagent kit. These were performed on the AdvanCETM FS96 system according to manufacturer’s guidelines (advancedanalytical, IA, USA). The amplified ORF was also re-sequenced by Sanger sequencing using primers listed in **Supplementary Table 1**.

#### Brachypodium distachyon

Mutation detection screenings were performed in the TILLING collection of chemically induced Brachypodium mutants, described in ^39^. TILLING by NGS consists to sequence 500 bp PCR fragments libraries prepared from 2600 individual genomic DNA pooled in two dimensions. A dual indexing system, one placed on the 5’adaptater, and the second one on the 3’adaptater, added by a two-step PCR (for primer sequence; see **Supplementary Table 1**) allow a direct identification of the sequence identities. The first PCR amplification is a standard PCR with target-specific primers carrying Illumina’s tail (**Supplementary Table 1**) and 10 ng of Brachypodium genomic DNA. Two microliters of the first PCR product served as a template for the second PCR amplification, with a combination of Illumina indexed primers (**Supplementary Table 1**). The sequencing step of PCR fragments was done on an Illumina Miseq personal sequencer using the MiSeq Reagent Kit v3 (Illumina^®^) followed by quality control processes for libraries using the PippinHT system from SAGE Sciences for libraries purification, and the Bioanalyzer™ system from Agilent®. To identify induced mutations, a bioinformatic pipeline, called “Sentinel” was used to analyze the data sequences (IDDN.FR.001.240004.000.R.P.2016.000.10000). Prediction of the impact of each mutation (**Supplementary Table 4**) was made with SIFT software as described in ^39^. The amplified ORF was also re-sequenced by Sanger sequencing using primers listed in **Supplementary Table 1**.

#### Sorghum

A pedigreed sorghum mutant library was established in the inbred line BTx623, which was used to produce the sorghum reference genome. This mutant library consists of 6,400 M4 grain pools derived from EMS-treated sorghum grains by single seed descent. Whole genome sequencing of a set of 256 lines uncovered 1.8 million canonical EMS-induced mutations ^39^. We searched the sorghum ortholog of the barley *COM1* in the aforementioned sequence database to identity plants carrying mutation. To confirm the mutations, the amplified ORF was also re-sequenced by Sanger sequencing using primers listed in Supplementary Table 1.

### Haplotype and network analysis

Genomic DNA from a core collection including 146 landrace and intermedium-spike barley accessions as well as 90 wild barley (Source data file for Supplementary Fig. 7) was PCR-amplified using specific primers to amplify full coding sequence of the barley *COM1* gene. Amplified fragments were used for direct PCR sequencing (Sanger method; BigDye Terminator v3.1 cycle sequencing kit; Applied Biosystems). A capillary-based ABI3730xl sequencing system (Applied Biosystems) at the sequencing facility of IPK was used to separate the fluorescently terminated extension products. Sequence assembly was performed using Sequencher 5.2.2.3. Visual inspection of sequence chromatograms was carried out using Sequencher to detect the corresponding SNPs. Network analysis of the nucleotide haplotypes was carried out using TCS v1.21 software (http://darwin.uvigo.es/software/tcs.html) ^43^.

### RNA extraction, sequencing and data analysis

#### RNA Extraction

For the RNA-seq study, immature spike tissues were collected from BW-NIL(*com1.a*) and Wt progenitor Bowman and the donor cv. Foma. Plants were grown under phytochamber conditions of 12h light (12 °C) and 12h dark (8 °C). Tissues were always collected at the same time slot (14:00 to 17:00) during the day at three different developmental stages including TM and GP, and pooled stages of LP+SP. Three biological replicated were applied that resulted in 27 individual tissue samples. The TRIzol method (Invitrogen) was applied to extract total RNA from immature spike tissues followed by removal of genomic DNA contamination using RNAse-free DNAse (Invitrogen). RNA integrity and quantities were analyzed via Agilent 2100 Bioanalyzer (Agilent Technologies) and Qubit (Invitrogen), respectively.

#### Preparation and sequencing of mRNA-Seq libraries

SENSE mRNA-Seq libraries (27 = 3 reps/3 stages /3 genotype) were prepared from 2 μg total RNA according to the protocol provided by the manufacturer (Lexogen GmbH, Vienna, Austria). Libraries were pooled in an equimolar manner and analysed electrophoretially using the Agilent 4200 TapeStation System (Agilent Technologies, Inc., Santa Clara, CA, USA). Quantification of libraries and sequencing (rapid run, paired-end sequencing, 2 × 100 cycles, on-board clustering) using the Illumina HiSeq2500 device (Illumina, San Diego, California, USA) were as described previously ^44^.

### Analysis of the RNAseq data

The reads from all three biological replicates were pooled per stage and each pool was independently mapped to barley pseudomolecules ^41^, (160404_barley_pseudomolecules_masked.fasta) using TopHAT2 ^45^. Gene expression was estimated as read counts for each gene locus with the help of featureCounts ^46^ using the gene annotation file Hv_IBSC_PGSB_r1_HighConf.gtf and fragment per million (FPM) values were extracted from the BWA-aligned reads using Salmon ^47^. Genes that showed FPM of 0 across all ^45^ samples were excluded from expression levels calculations. Expression levels were normalized by TMM method and *p*-values were calculated by an exact negative binomial test along with the gene-specific variations estimated by empirical Bayes method in edgeR ^48^. The Benjamini-Hochberg method was applied on the *p*-values to calculate *q*-values and to control the false discovery rate (FDR). Differentially expressed genes (DEGs) were defined as *q*-value < 0.05, log2 fold change > 1 or < −1.

### Quantitative RT-PCR

Tissue sampling, RNA extraction, qualification and quantification was performed as described above. Reverse transcription and cDNA synthesis were carried out using SuperScript III Reverse Transcriptase kit (Invitrogen). Real-time PCR was performed using QuantiTect SYBR green PCR kit (Qiagen) and the ABI prism 7900HT sequence detection system (Applied Biosystems). Each qRT-PCR comprised at least four technical replicates, and each sample was represented by three biological replicates. The *Actin* gene-based primers (**Supplementary Table 1**) were used as the reference sequence. qRT-PCR results were analyzed using SDS2.2 tool (Applied Biosystems) in which the presence of a unique PCR product was verified by dissociation analysis. Significance values were calculated using *Student’s t*-test (two-tailed). The relevant primer sequences per species are detailed in **Supplementary Table 1**.

### Phylogenetic analysis

A comprehensive analysis of TCP proteins in grasses was already available we therefore focused only on constructing a detailed phylogeny of the COM1 protein among grasses and the barley TCP genes. Thus, barley COM1 was then queried against Ensembl Plants database to retrieve its orthologs or homologs from other grasses. The same database was also used to extract all barley TCP proteins. In case of COM1, protein and DNA sequence of the paralog and homologous genes from each of the grass species were retrieved. To re-check their homology with barley COM1, the retrieved sequences were blasted back against the barley genome. For phylogenetic analysis, protein sequences were initially aligned using the algorithm implemented in CLC sequence viewer V7.8.1 (https://www.qiagenbioinformatics.com). UPGMA tree construction method and the distance measure of Jukes-Cantor were implemented for constructing the phylogenetic tree using CLC sequence viewer. The bootstrap consensus tree inferred from 1000 replicates was taken to represent the evolutionary relationship of the sequences analyzed.

### mRNA in situ hybridization

In case of *COM1*, three separated segments (excluding the TCP domain) from the *COM1* gene each containing 300-360 bp were synthesized (probe 1 and 2, GenScript Biotech, Netherlands, Source data files) or amplified (probe 3) using cDNAs isolated from immature spikes of cv. Bonus and specific primers (Supplementary Table 1). The resulting products were cloned into pBluescript II KS (+) vector (Stratagene, La Jolla, CA, USA and GenScript Biotech, Netherlands). Linearized clones by HindIII or NotI were used as templates to generate antisense (HindIII) and sense (NotI) probes using T3 or T7 RNA polymerase. *In situ* hybridization was conducted with a single pool of the three aforementioned probes as described previously ^49^.

For the *HvLG1* gene, a single probe, derived from the third exon (see Source data files for probe sequence) was synthesized (GenScript Biotech, Netherlands). The aforementioned approach described for *COM1* was conducted for *in situ* hybridization.

### Scanning electron microscopy

Scanning electron microscopy (SEM) was performed on immature spike tissues at five stages including triple mound, glume, lemma, stamen, and awn primordium from greenhouse-grown plants. SEM was conducted as described elsewhere ^50^.

### DNA preparation

DNA was extracted from leaf samples at the seedling as described before ^13^. Plants for which the DNA was prepared included barley, *Sorghum* and *Barchypodium*. That included either mapping population, TILLING mutants or both.

### Palea anatomical and TEM analyses

For anatomical study as well as transmission electron microscopy (TEM), plant material consisting of intact spikes was collected shortly before anthesis. Spikelets containing no grains were used for dissecting paleae that were subsequently stored in fixative (4% FA, 1% GA in 50 mM phosphate buffer). Central spikelets (in case of barley) were isolated and placed in a 15 ml test tube containing 10 ml fixative, followed by extensive degassing until all probes had settled. Material was stored in a fridge until use. After three washes with A.D., lemma and palea were isolated by cutting away a small part at the base of the spikelet. Isolated paleae were placed in a flat bottomed mold filled with 4% liquid agarose (~60°C). After setting, agarose blocks were removed from the mold and the encapsuled Palea was cut into 1-2mm wide sections using fresh razor blades. The embedding in agarose facilitated the cuttings while preventing unnecessary damage to the probes. After embedding in Spurr resin (see next page) semithin sections of 2 μm were cut on an Leica Ultracut. Sections were allowed to be baked in a droplet of 0,02% Methylene blue/Azur blue on a heating plate set at 90°C. Recordings were made using a Keyence VHX-5000 digital microscope (Keyence Germany GmbH, Neu-Isenburg, Germany).

### Sequence information and analysis

Unpublished sequence information for the BAC contigs 44150 spanning the interval between NGS084 and NGS094) was made available from the international barley sequencing consortium (through Nils Stein). This sequence information was used for marker development during high resolution mapping, map-based cloning and *COM1* gene identification. Later on, the initial contigs 44150 sequence information was re-checked and confirmed with the high-quality barley genome assembly and annotation data ^27^.

### Whole genome shotgun sequencing of BW-NIL(*com1.a)*

A whole-genome shotgun library was constructed using standard procedures (TruSeq DNA; Illumina) and quantified using real-time PCR. Cluster formation using the cBot device and paired-end sequencing (HiSeq2000, 2 × 101 cycles) were performed according to the manufacturer’s instructions (Illumina).

## Acknowledgments

We thank the US Department of Agriculture–Agricultural Research Service (USDA-ARS), the National Small Grains Collection, Aberdeen (ID); the Nordic Genetic Resources Center (NordGen), Alnarp, Sweden; and the IPK Genebank, Germany, for providing the mutants and germplasm for haplotype analysis. The authors would like to thank Dr. Shun Sakuma for fruitful discussions and help in conducting mRNA *in-situ* hybridizations. We are thankful to Anne Fiebig for help with data submission to ENA and Sandra Driesslein, Jenny Knibbiche, Mechthild Pürschel, Ines Walde, Kerstin Wolf, Marion Benecke and Kirsten Hoffie for excellent technical assistance.

## Author contributions

T.S. conceived the idea for the study, designed and monitored experiments, and analyzed data. N.P. expanded the idea for the study, designed and performed experiments and analyzed data; C.T. executed the mRNA *in-situ* hybridizations. M.M. conducted microscopic analyses of cellular structures in paleae. T.N. analyzed RNA-seq data. U.L. provided irregular and intermedium barley spike mutants. T.R. executed SEM analyses. T.Schm. conducted sequence read mapping to unpublished barley genomic sequences for SNP calling. R.B., A.H. and L.A. performed the initial whole-genome shotgun sequencing of the parental genotypes for mapping. R.K. was involved in the phenotypic analysis of *com1.a* and RT-qPCR analyses of *COM1* in barley *vrs4* mutant. H.M.Y. provided sequences from a barley diversity panel for haplotype analysis and was involved in the RT-qPCR analysis. R.S., M.D. and A.B. provided the Brachypodium TILLING resource; N.S. provided the barley TILLING resource; Z.X. provided the sorghum TILLING resource. N.P. and T.S. wrote the manuscript including contributions from co-authors. All authors have seen and agreed upon the final version of the manuscript.

## Data and materials availability

Barley mutants are available from TS under a material transfer agreement (MTA) with IPK-Gatersleben. All data are available in the main text or online materials. The RNA-seq data and the whole genome shotgun (WGS) sequences of *com1.a* mutant have been submitted to the European Nucleotide Archive under accession number PRJEB35746 and PRJEB35761, respectively. COM1 sequences are available with the corresponding ID mentioned in the current study in the public databases https://plants.ensembl.org/ & https://apex.ipk-gatersleben.de/apex/f?p=284:10 and are in the process of submission to NCBI as well. The source data underlying figures (Fig. 5Q-S, Fig. 6D-G, Fig. 7I-N, Fig. 8A-B, Fig. 8C-D, and Supplementary Fig. 7) and tables (Supplementary Table 5) are provided as Source Data files.

