## Supplementary material for "COMPOSITUM 1 (COM1) contributes to the architectural simplification of barley inflorescence via meristem identity signals": COM1_Suppl (from another version of this preprint: version 1, 2020-02-19)

### Supplementary Information for

**Title:**

**COMPOSITUM 1 (COM1) contributes to the architectural simplification of barley inflorescence via cell wall-mediated and meristem identity signals**

**Authors:**

N. Poursarebani[1], C. Trautewig[1], M. Melzer[1], T. Nussbaumer[3,4], U. Lundqvist[5], T. Rutten[1], T. Schmutzer[1,2], R. Brandt[1], A. Himmelbach[1], L. Altschmied[1], R. Koppolu[1], H. M. Youssef[1,6], N. Stein[1], Z. Xin[7], T. Schnurbusch[1,2]

**This file includes:**

**Supplementary Notes**

**References**

**Supplementary Figs. 1 to 12**

**Supplementary Tables 1 to 6**

**Captions for Supplementary Source Data 1**

#### Supplementary Notes

##### *COM1* positional cloning in barley

The genetic map was conducted by screening ~6,000 gametes for recombination events in an $F_2$ population (Bowman x *com1.a)*. After which, fifteen critical recombinant $F_2$-derived $F_3$ families (i.e., 16 plants per family; **Supplementary Table 2- 3**) were further analyzed that unambiguously confirmed the genetic interval detected. Resequencing a set of 20 barley spike-branching mutants, using both CDS and promoter specific primer pairs (**Supplementary Table 1 and S4**), revealed that five of them, i.e. *Mut.3906*, *int-h.42, int-h.43* and *int-h.44*, and *com1.i*, lost the same transcription factor as found missing in the *com1.a* mutant (**Supplementary Fig. 3, Supplementary Table 1 and 4**). All five mutants also showed the flat-palea phenotype observed in the mutant *com1.a* (**Supplementary Fig. 3**). Allelism tests of *com1.a* with *Mut.3906* indicated that they are allelic to each other. Furthermore, we PCR-screened a barley TILLING populations from cv. Barke (two-rowed) for the CDS of the candidate gene. Four homozygous M3 plants (M3.15104, M3.4406, M3.13729 and M3. 2598) carrying SNP mutations inside the DNA binding domain as well as two heterozygous M3 lines M3.4063 and M3.9299 with SNP mutation outside the domain were identified (**Fig. 2B**). All six SNP mutations caused amino acid substitution in conserved positions (**Fig. 2B**). They all transmitted a branched spike as was revealed by the phenotypes of the corresponding M4 and M5 homozygous plants (**Fig. 2B; Supplementary Fig. 4–5; Supplementary Table 4**). Interestingly, from the six TILLING mutants, only two, with mutation within the TCP domain, showed either a true flat-palea phenotype with a complete loss of the infolding (line 2598, exhibiting also the most severe branching), or only a mild change in the palea shape (line 4406) (**Supplementary Fig. 4**). Thus, penetrance of the mutant flat-palea phenotype depended on the type and position of the amino acids substitution

(**Supplementary Fig. 4**, legend). Taken together, these data indicate that the mutations in this transcription factor caused the two *com1* phenotypes.

**Development and characterized of BW-NIL (*com1.a com2.g*) double mutants**

20 BW-NIL(*com1.a com2.g*) double mutants homozygote plants along with 20 plants from each of the single mutants and 20 wild type cv. Bowman plants, were grown and used for phenotyping. The DM plants out-performed either single mutants in supernumerary spikelet and branch production and grain number per spike, which was elevated by ~50% in DMs as compared to single mutants alone (**Supplementary Fig. 6 and 12**, legend). Notably, all tillers from DMs showed both supernumerary spikelet formation and spike-branching phenotypes. In addition to the ramification observed in central SMs, SEM analysis of DM plants also showed floral reversion of the lateral SMs (**Supplementary Fig. 6**). Both events were highly penetrant and pronounced in the DM's immature spikes that led 100% of the plant tillers to exhibit a DM-specific enhanced spike-branching phenotype at maturity (**Supplementary Fig. 6 and 12**). The uncommon enlargement of the glum primordia (Blue asterisk in Fig. 1O) was seen only in ~5% of the tillers in single mutant *com1.a* whereas 100 % of the tillers in *com1.acom2.g* double mutants showed this enlargement (**Supplementary Fig. 12**).

**TILLING analysis of *BdCOM1* in *Brachypodium***

Our TILLING analysis in *Brachypodium distachyon* revealed several mutations in the *BdCOM1* homolog from which two (M4. 5446; Q116* and M4. 8373; S146N) with predicted severe damages to the protein domain and function were phenotypically characterized (**Supplementary Table 4**). Hence, per M4 plants, only homozygous M5 plants either with mutant genotype aa (3 to 4 plants) or wild type bb (3 to 4 plants) were selected. Per M5 plants, 10 M6 plants were grown. The homozygote states of the corresponding M5 and M6 mutant family transmitted a defect in palea structure (**Fig. 3**). In which, we always observed a palea scissor-like structure in all florets in mutants, as it collapses easily due to external mechanical pressure. In contrast, palea in Wt never made such structure while applying same external hand-pressure. Histological analyses of the *Brachypodium* mutants' palea revealed no obvious change in cell expansion (**Fig. 4**).

#### Supplementary figures

**Supplementary Figure 1. Creation of induced Bowman near isogenic line (NIL), mapping population, marker resource and the low-resolution linkage map of *com1.a* in barley.** (**A and B**) The *com1.a* phenotype was introduced (after its induction in barley cv. Foma) into a two-rowed barley cv. Bowman by six time backcrosses as reported previously [1]. Green area in chromosome 5H corresponds to the introgressed genomic segment caring the underlying *com1.a* mutation. (**C**) The resulted BW near isogenic line (NIL) of *com1.a* allele; BW-NIL (*com1.a*) was crossed to cv. Bowman to generate $F_2$ population used in genetic mapping. (**D**) BW-NIL (*com1.a*) was whole genome shotgun (WGS) sequenced to 10x coverage (Online Materials) and compared with already published WGS of Bowman [2]. (**E**) Represents alignment of the BW-NIL (*com1.a*) sequence assembly against the physically localized Bowman WGS sequence contigs. The corresponding SNPs (that were used for marker development) derived from the 5H introgression segment are plotted in the outer circle. (**F**) Genetic linkage mapping of *com1.a* in barley, derived from the $F_2$ mapping population (see part C). SNPs derived from E located within the introgressed region, were used for marker development. Markers in red selected for high-resolution genetic mapping. A-B were performed and published previously [1]. C-F are entirely performed in the present study, except WGS of the wild type cv. Bowman.

**Supplementary Figure 2: TEM based cell wall structure in wild type palea versus mutant *com1.a*.**

**Left-ide**; cross section of wild type palea in which different cell positions (used to image the cell walls shown in right panel) across cell layers (see Fig. 1S-T for the layer IDs) are labeled. **Right-side**; Cell walls thickness in Wt is depicted compared with that of mutant for each position of A-I; labeled accordingly in Left panel. BW-NIL (*com2.g*) mutant, that is visually similar in palea phenotype/structure with that of Wt cv. Bowman, were used here to compare with the mutant *com1.a*.

**Supplementary Figure 3. Additional mutant alleles of spike branching in barley.** Different *com1* induced mutant alleles identified by resequencing of primers correspond to CDS and potential promotor region of *COM1*. The corresponding palea is shown in the upper –right side of each spike image. See also **Supplementary Table** 1 and 4.

**Supplementary Figure 4. Phenotype and COM1 protein sequence alignment in barley TILLING lines.** (**A**) A representative display of branch formation of the six barley TILLING mutant plants derived from barley cv. Barke. The corresponding palea is shown in the upper – right side of each spike image. (**B**) Protein sequence alignment of the six mutants, some located within the corresponding TCP domain (the red box). Mutation pointed with red arrows show severe (dark red; M4.2598) and very mild (light red; M4.4406) palea phenotypes, while green arrows show mutation with no palea phenotype.

**Supplementary Figure 5. Tillering related characters in barley TILLING lines. (A)** Average tiller number and (**B**) spike number per plant of the three spike-branching barley TILLING lines are compared against wild type cv. Barke. This was performed to check whether *COM1* expression detected in tiller buds (**Fig. 4B**) play any role in tiller formation.

Twenty plants were grown per genotype under greenhouse conditions. *P* values were determined using the Student's *t* test.

**Supplementary Figure 6. Immature spike and yield related components in double mutant BW-NIL (com1.a/com2.g) (DM).** (A) Lateral view of a DM immature spike at late GP stage. (**B-D**) Close-up and complete view of a DM immature spike; depict reversion of lateral and central SMs as well as glume primordium into the IM-like meristems. (**E-J**) Yield components of the DM plants, and the corresponding single mutants com1.a and com2.g in comparisons to the Wt cv. Bowman. Data are based on a single greenhouse-condition experiment and on averages of 20 plants (390 to 540 spikes) per phenotypic class.

**Supplementary Figure 7. Phylogeny of barley TCP proteins and the phylogeny-based model of *COM1* evolution. (A)** 1000 bootstraps based UPGMA phylogenetic tree of 22 barley TCP proteins. (**B).** Depicts origin and divergence of two (out-) paralogs of REP1/COM1 (α) and MSD1 (β*;* [3] clades from a common ancestor [4]. This divergence seems to happen after genome duplication in grass common ancestor and before the grass speciation (e.g. the separation of *Triticeae* from *Non-Triticeae* species). More importantly, in case of paralogs α clade; although the clade members showed no duplication (no in-paralogy) after speciation, except in maize (Fig. 2C; light-grey block), they have gone through a functional modification that separate *Non-Triticeae* function (α1) from that of *Triticeae* subclade (α2). Which, coincidentally, the respective modification separates inflorescence shapes between the two groups of grass plants by regulating contrasting phenotypes of formation versus inhibition of branches. Whether out-paralog β also differs in function among grass (to be as β1 and β2) remains to be investigated. So far, only functional characterization of β in sorghum, the *SbMsdI,* has been reported [3]. Lines in brown represent the speciation process.

**Supplementary Figure 8. Amino acid sequence alignment of the COM1 homologs among grass**. The green box shows the TCP domain conservation.

**Supplementary Figure 9. Network analysis of *COM1*.** Depicts grouping of 237 barley accessions (including 90 wild barleys) producing 12 haplotypes; comprising two main haplotypes: namely Hap1 (154 wild and landraces) and Hap2 (64 wild and landraces) with Hap1 being assigned as ancestral. The remaining 10 haplotypes, except Hap8 that contains two landraces comprised only wild barleys (16 accessions), independently raised from Hap1 during the course of evolution. Accessions representing each of the 12 haplotypes were grown and carefully inspected for the spike and palea phenotype for which no obvious phenotypic alteration was observed. Sequences were obtained using primers for the CDS region.

**Supplementary Figure 10. Branch formation in *vrs4* mutant (*mul1.a*).** (**A**) Mature spike of wild type progenitor cv. Montcalm with determinate triple spikelet meristem. (**B-D**) Mature spikes of *vrs4* mutant MC (*mul1.a*) showing various levels of branch proliferation at the spike base and middle portion of the spike. See also [5]. *COM1* **(E)** and *COM2* **(F)** transcripts in BW-NIL(*vrs4.k*) (red) and *com1.a* (green) mutants, respectively. Mean ±SE of three biological replicates. *P* values were determined using the *Student's t* test.

**Supplementary Figure 11. Transcriptome analysis of *com1.a* using RNA seq.** Heat map of all DE genes (found in RNA seq; n = 3 reps/3 stages /3 genotype) conjointly in the BW-NL (*com2.g*) as compared to the corresponding wild type cv. Bowman and cv. Foma. The scale bar at the top of the heat map indicates the transcript level of differentially regulated genes observed between wild type and mutant; blue color indicates down-regulation while red shows up-regulation. Of the highly upregulated genes in mutant *com1.a* was HORVU3Hr1G055600; NAD-dependent epimerase/dehydratase gene family; associated with increased growth [6]. Other highly upregulated genes involved in plant reproduction (HORVU7Hr1G054320;, type I MADS-box transcription factor family protein; [7,8], as well as the floral meristem determinacy (HORVU7Hr1G018780; Homeobox-leucine zipper protein family protein 4; homology to HAT1 gene in *Arabidopsis thaliana*) [9]. Up regulation of these gene sets is fairly in line with the observed spike-branching phenotype of the mutant *com1.a*. See the main text; for illustration of the relevant genes that are downregulated in mutant.

**Supplementary Figure 12. The extended model of *COM1* regulations.** (**A**) Model of Wt *COM1* transcriptional regulation based on "down-regulation" of the Wt allele (**B**) The resulted spike phenotype (% of tiller per mutant plant) due to the respective gene(s) loss-of-funation. Data are based on a single greenhouse-condition experiment and on averages of 20 plants (390 to 540 spikes) per mutant class (*com1.a*, *com2.g* and the respective double mutants).

#### Supplementary figures

**Supplementary Figure 1.**

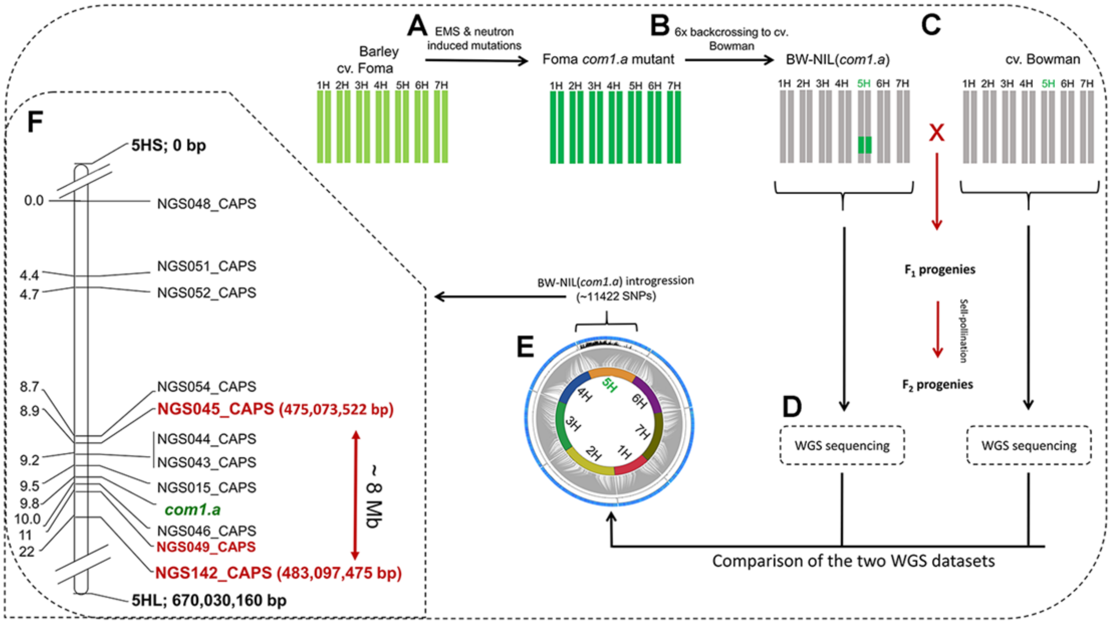

A
EMS & neutron induced mutations
Barley cv. Foma
1H 2H 3H 4H 5H 6H 7H
B
6x backcrossing to cv. Bowman
Foma com1.a mutant
C
BW-NIL(com1.a)
cv. Bowman
X
F1 progenies
Self-pollination
F2 progenies
D
WGS sequencing
WGS sequencing
Comparison of the two WGS datasets
E
BW-NIL(com1.a) introgression (~11422 SNPs)
5H
4H
6H
3H
7H
2H
1H
F
5HS; 0 bp
0.0 NGS048_CAPS
4.4 NGS051_CAPS
4.7 NGS052_CAPS
8.7 NGS054_CAPS
8.9 NGS045_CAPS (475,073,522 bp)
NGS044_CAPS
9.2 NGS043_CAPS
9.5 NGS015_CAPS
9.8 com1.a
10.0 NGS046_CAPS
11 NGS049_CAPS
22 NGS142_CAPS (483,097,475 bp)
~ 8 Mb
5HL; 670,030,160 bp

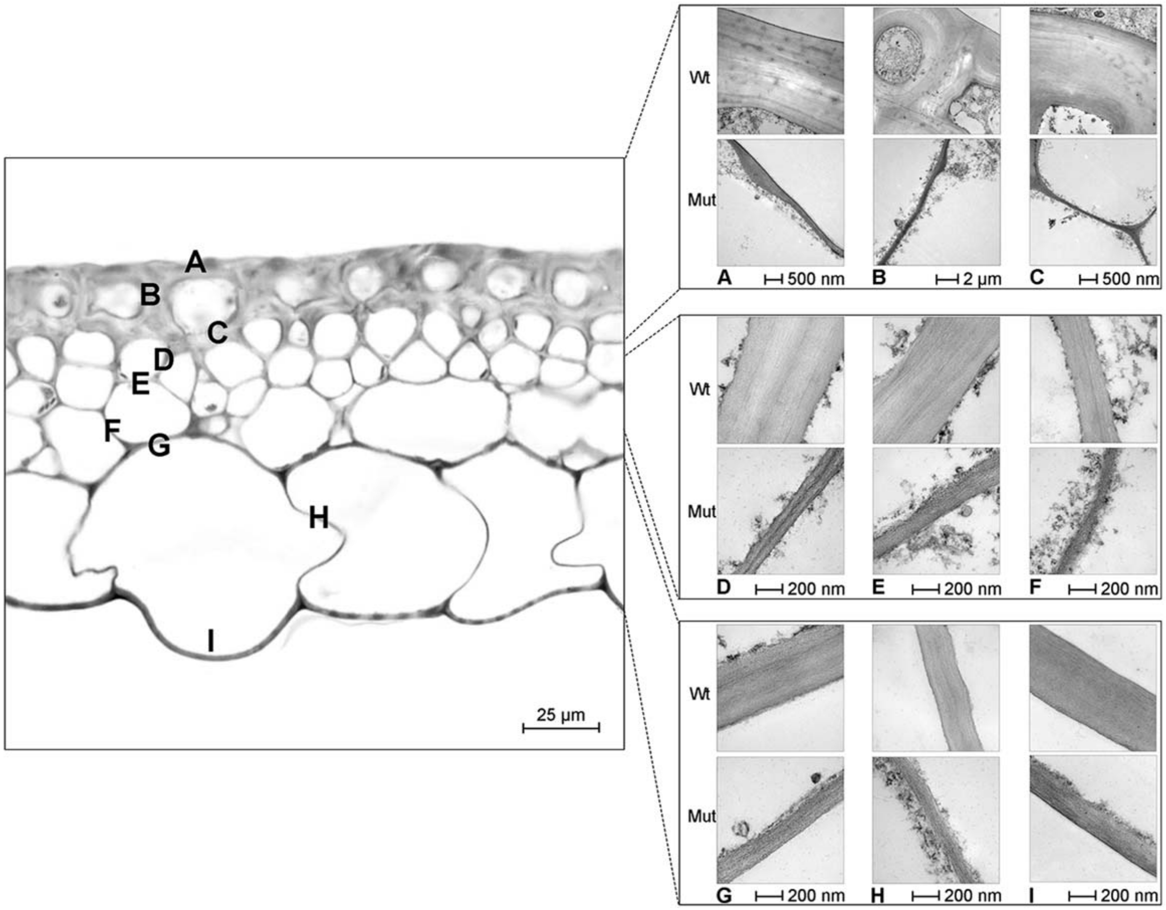

**Supplementary Figure 2**

**Supplementary Figure 3**

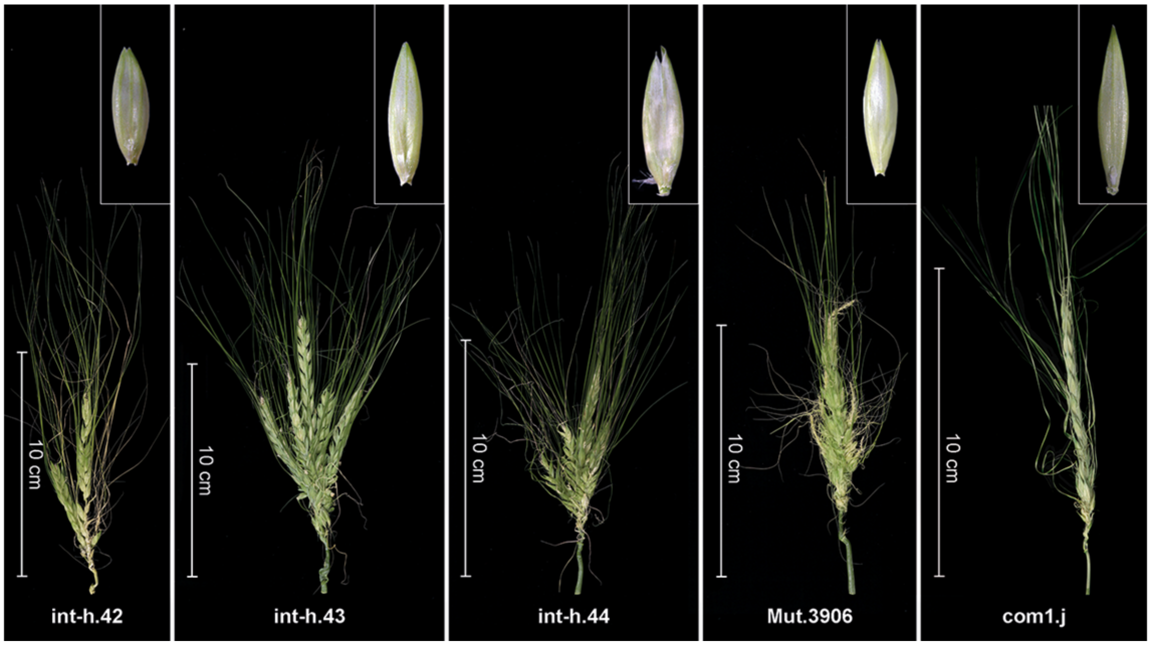

#### Supplementary Figure 4

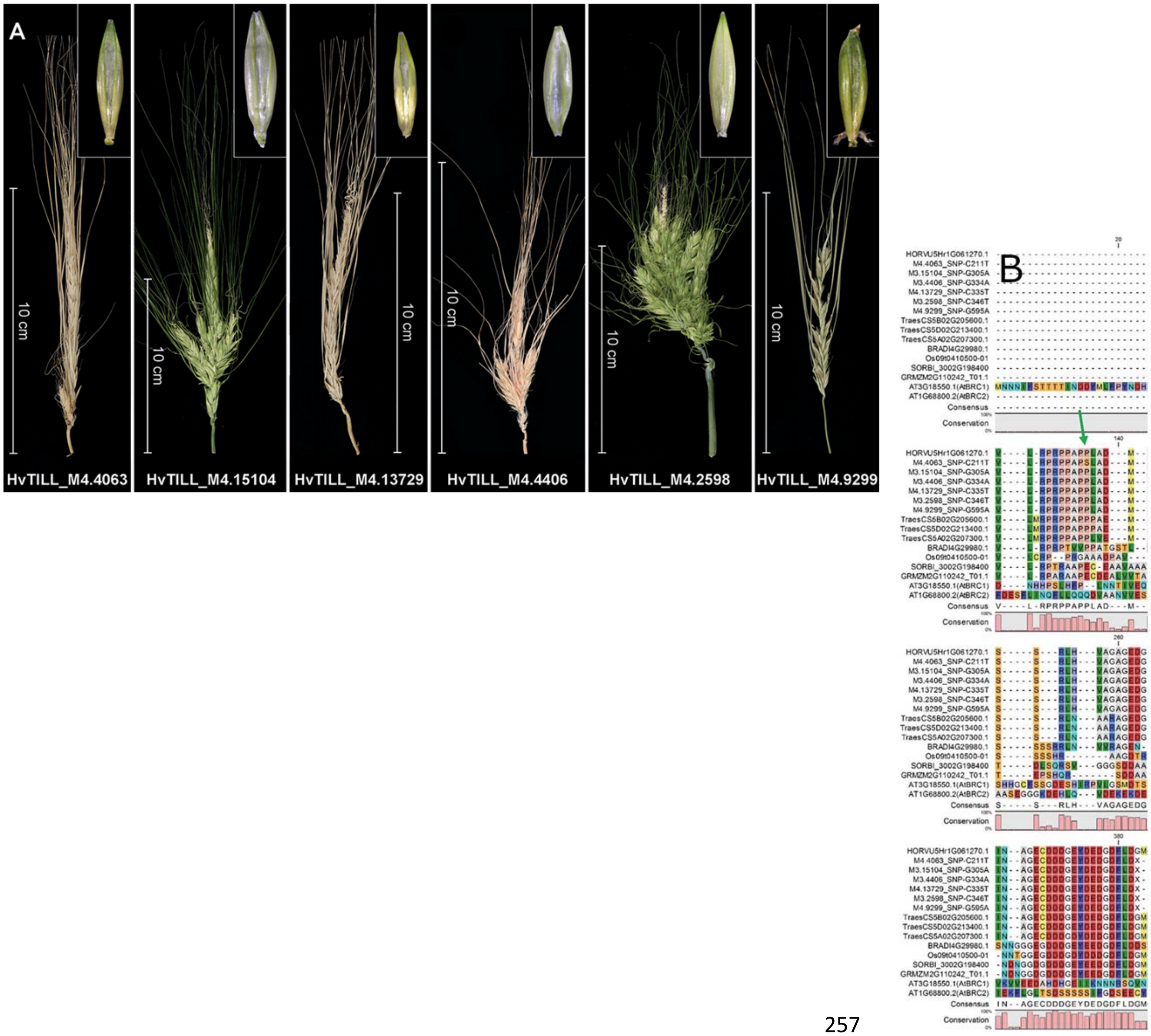

#### Supplementary Figure 5.

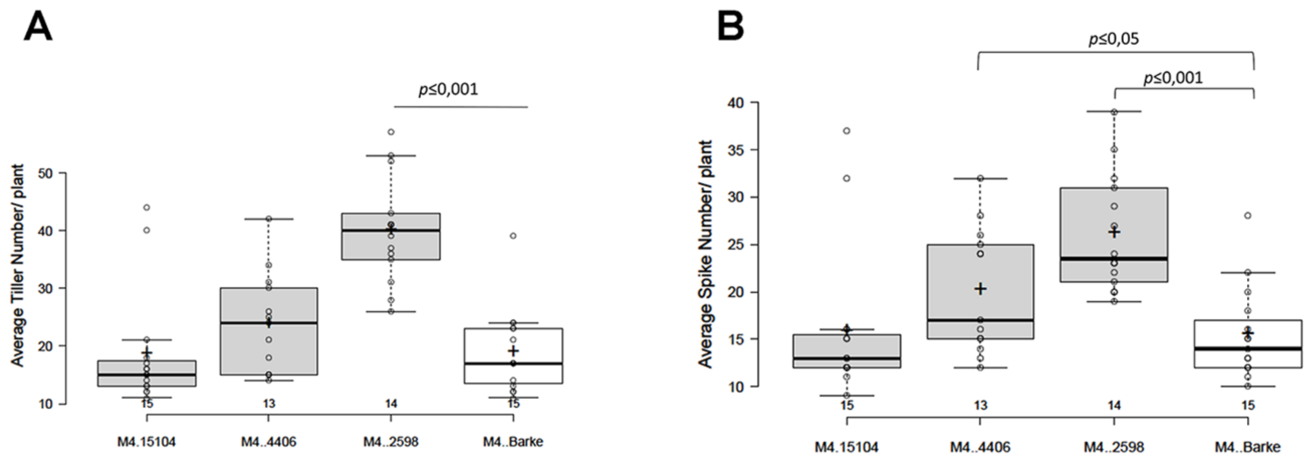

#### Supplementary Figure 6

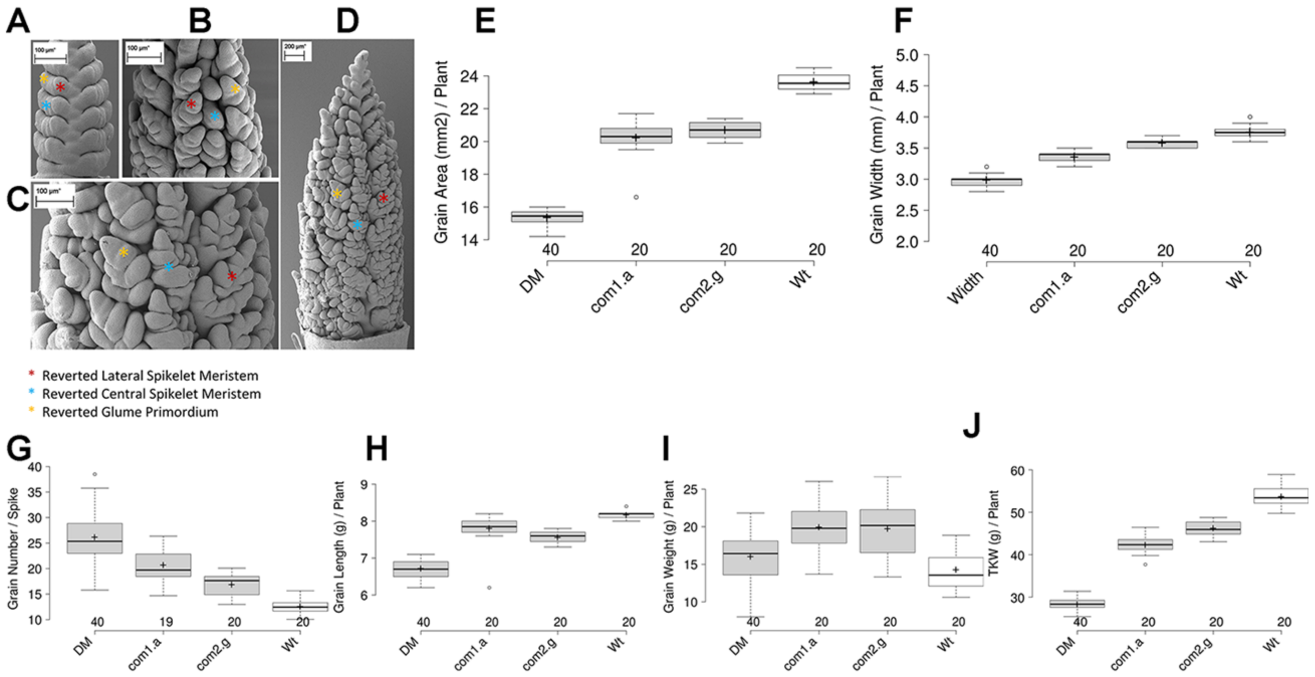

.

#### Supplementary Figure 7

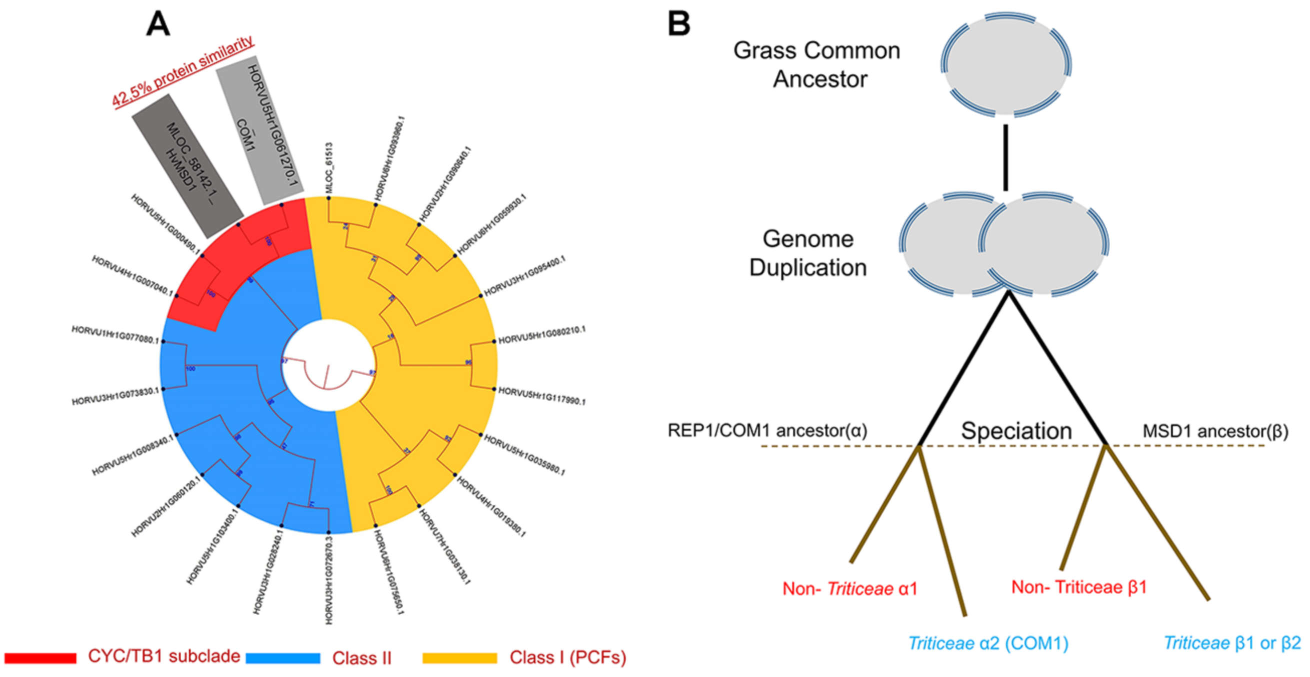

**Supplementary Figure 8**

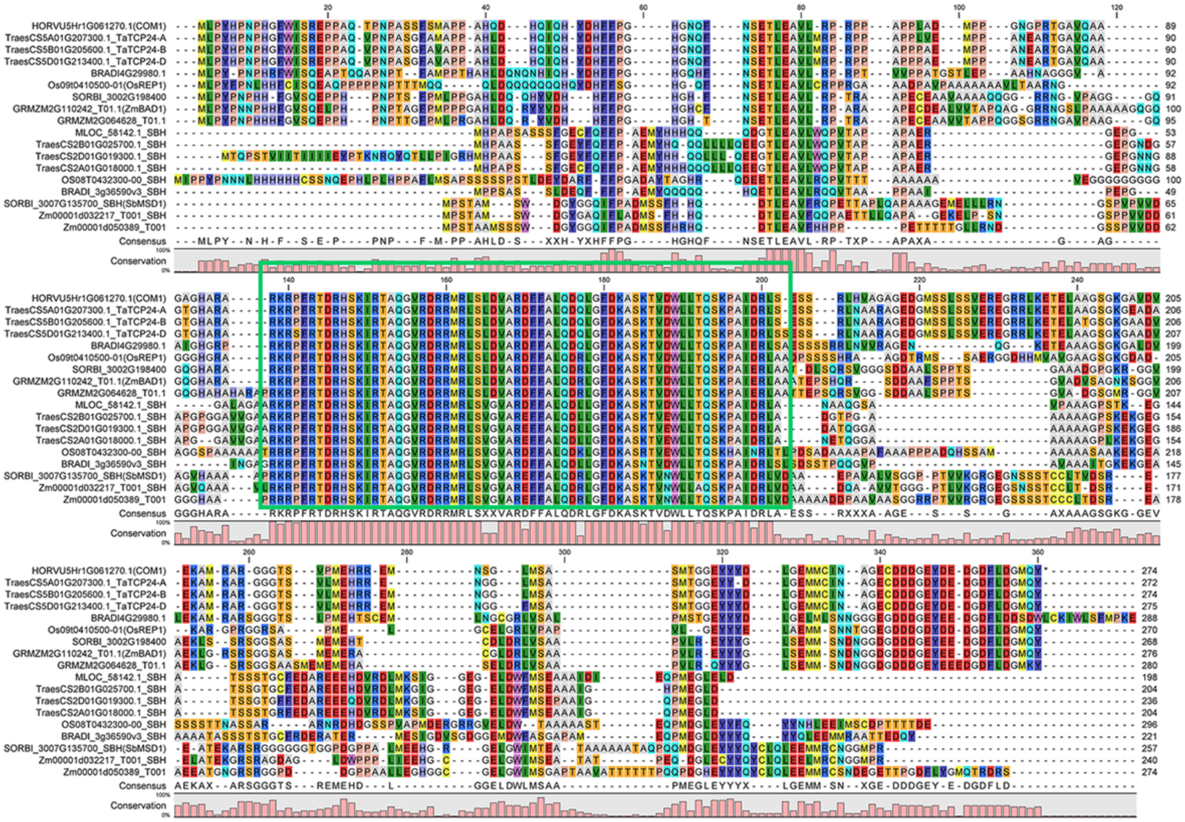

#### Supplementary Figure 9

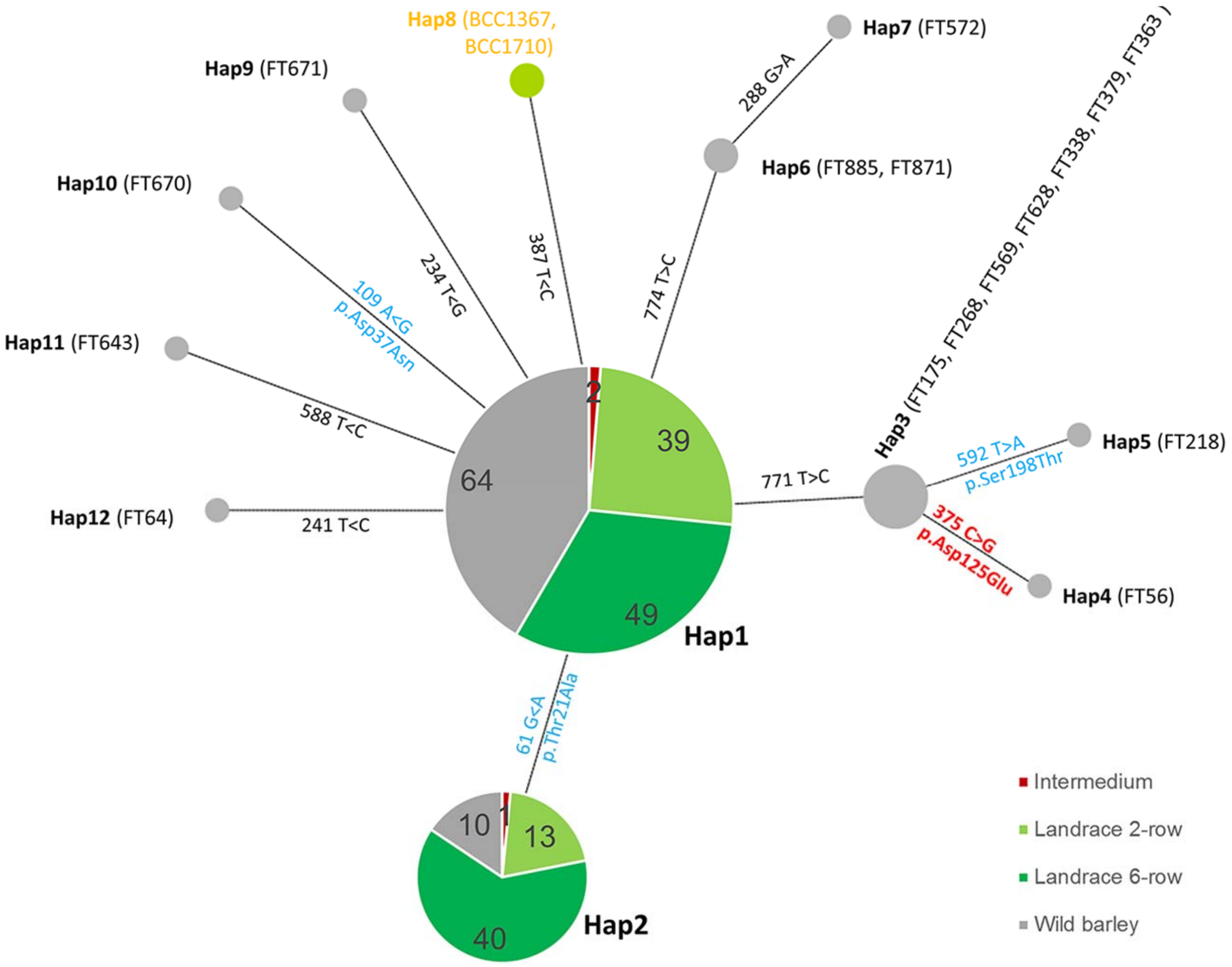

**Supplementary Figure 10**

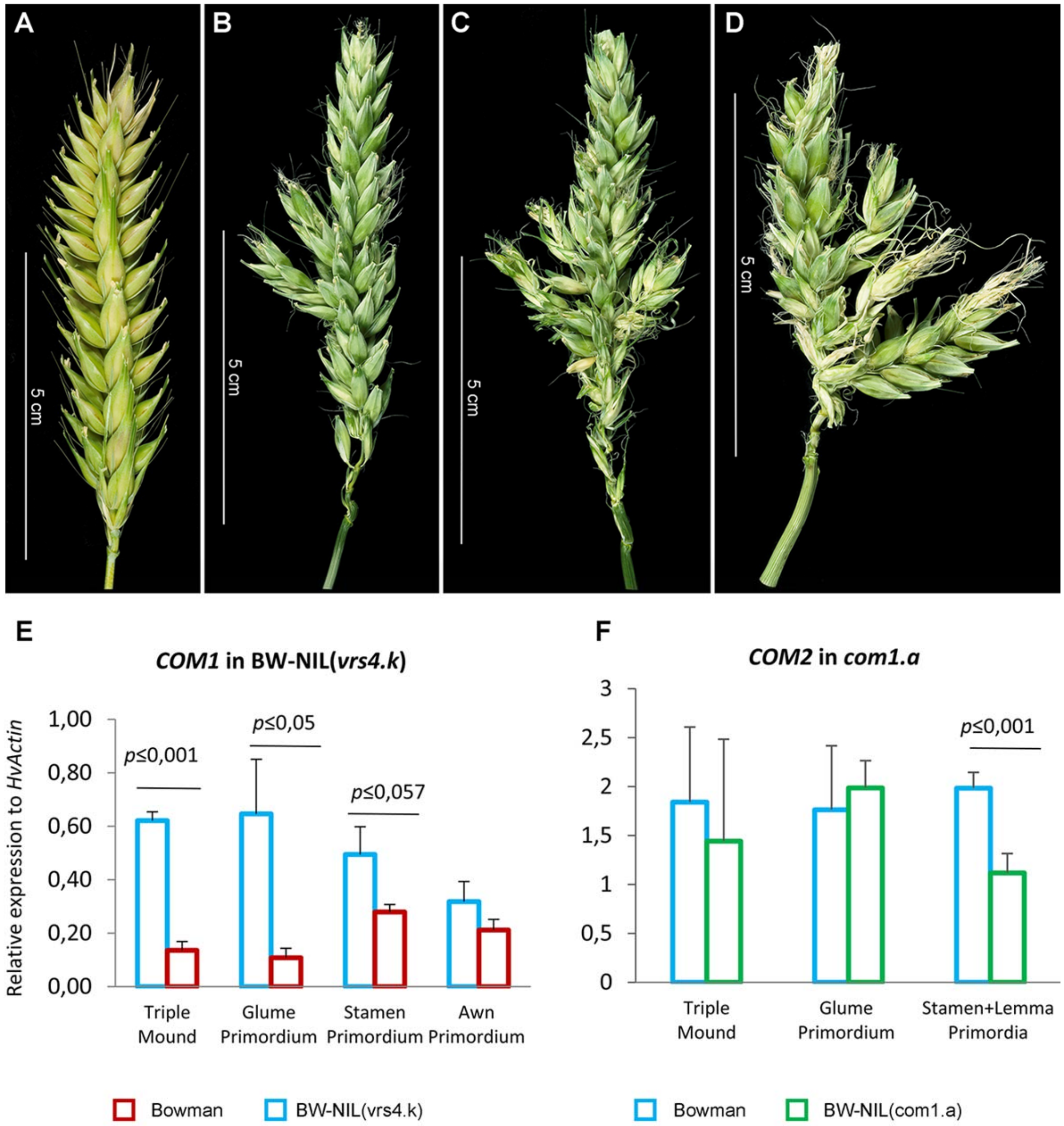

**Supplementary Figure 11**

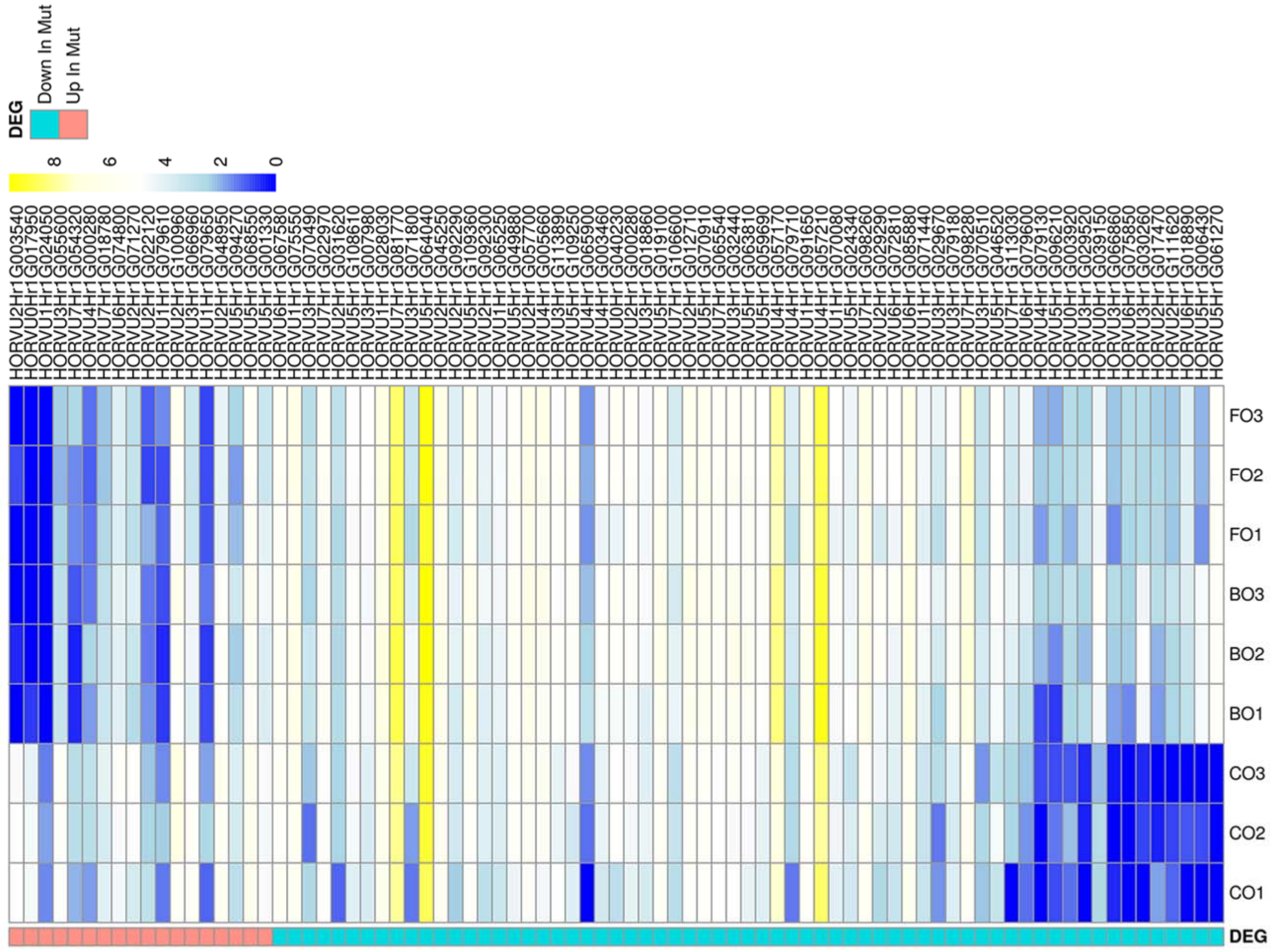

**Supplementary Figure 12**

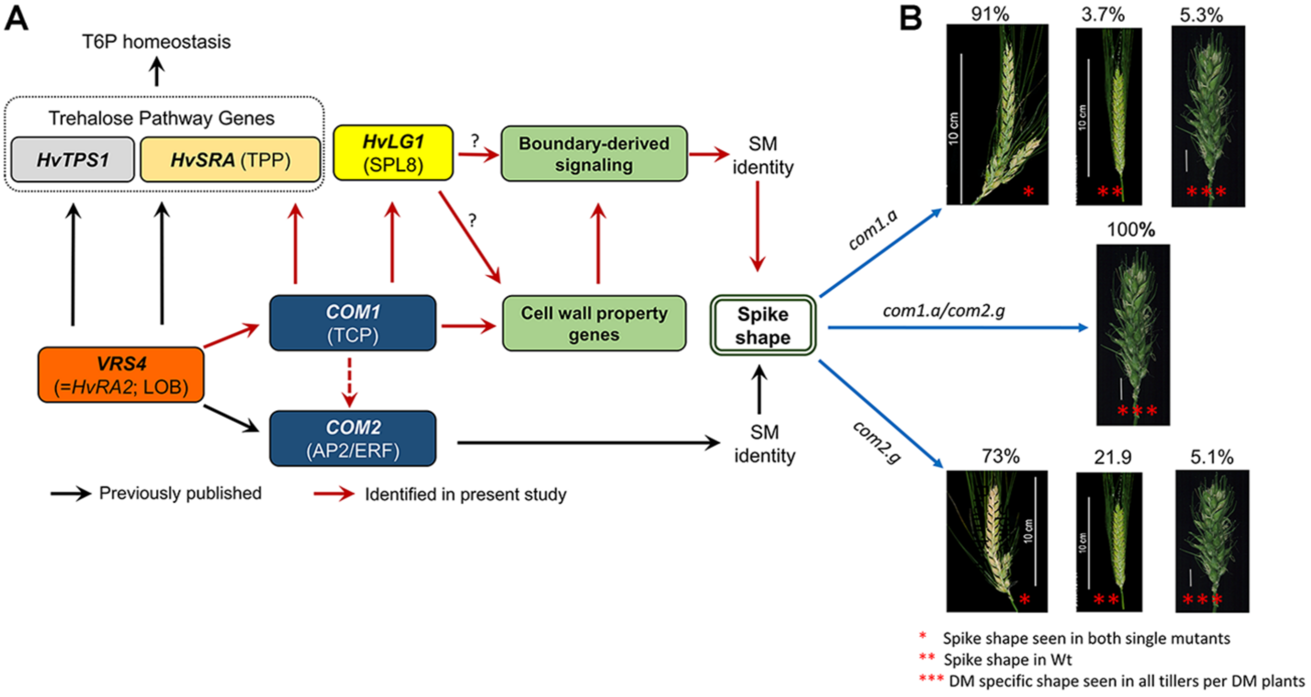

#### Supplementary tables

**Supplementary Table 1. List of the primers**

**Supplementary Table 2. Graphical genotyping of the critical $F_2$ recombinants used to develop $F_3$ families**

**Supplementary Table 3. Genotypic and phenotypic data of the $F_3$ progenies**

**Supplementary Table 4. List of the TILLING as well as induced mutants per corresponding species**

**Supplementary Table 5. Phenotypic data of the sorghum-TILLING mutants**

**Supplementary Table 6. Haplotypes identified for *COM1* in barley**

| Supplementary Table 1. List of the primers | | | | | | | |
|---|---|---|---|---|---|---|---|
| Primer ID | Species | Application | Marker Type | Orientation | tm | Seq | RE |
| NGS0015 | Barley | genetic mapping | CAPS | FORWARD | 60,36 | ACTACAGGAGTGCTGCTGGTAAA | SfaNI |
| NGS0015 | Barley | genetic mapping | CAPS | REVERSE | 62,07 | TTGCGGTATGCAACTCTCAACT | |
| NGS043 | Barley | genetic mapping | CAPS | FORWARD | 61,56 | TCGAGACTGAGGTAGTGGGACTT | PstI |
| NGS043 | Barley | genetic mapping | CAPS | REVERSE | 62,06 | CCGAAGGTGGTCAATAGACAAAG | |
| NGS044 | Barley | genetic mapping | CAPS | FORWARD | 61,43 | GCAACTGGGATTCGATCTCTTAG | EcoRV |
| NGS044 | Barley | genetic mapping | CAPS | REVERSE | 61,14 | CTAAAGCCTTGCACAAAGTTGG | |
| NGS045 | Barley | genetic mapping | CAPS | FORWARD | 62,69 | ACACCGAGATGTTGTTGGAAGAG | BtgI |
| NGS045 | Barley | genetic mapping | CAPS | REVERSE | 62,07 | ATGGATACGGAAGCCAGTGTCTA | |
| NGS046 | Barley | genetic mapping | CAPS | FORWARD | 62,15 | GATACACTTAAGGCCAAACGGTTC | HinfI |
| NGS046 | Barley | genetic mapping | CAPS | REVERSE | 61,58 | TACGTCAGCTGGACACACACATA | |
| NGS048 | Barley | genetic mapping | CAPS | FORWARD | 62,03 | CTCCTACGTGATTCACTGTGTCG | SspI |
| NGS048 | Barley | genetic mapping | CAPS | REVERSE | 61,62 | TTCAGAGGCTGAAGAAAGAGAGC | |
| NGS049 | Barley | genetic mapping | CAPS | FORWARD | 61,95 | GGTGATAAATCCACTCCAGCAAC | BbsI |
| NGS049 | Barley | genetic mapping | CAPS | REVERSE | 61,82 | GTCAAAGTGGAGAAGCTGCAAA | |
| NGS051 | Barley | genetic mapping | CAPS | FORWARD | 61,97 | TGGTCGTTGGCTTCTCTAGTTTC | PstI |
| NGS051 | Barley | genetic mapping | CAPS | REVE | 62, | GAACGAAATCAACACAGGAGA | |

| | | | | | | | |
|---|---|---|---|---|---|---|---|
| | | | | RSE | 17 | CAC | |
| NGS052 | Barley | genetic mapping | CAPS | FORWARD | 61,95 | GAGAGTAGGCAGATCCAACGAAA | TagI |
| NGS052 | Barley | genetic mapping | CAPS | REVERSE | 61,96 | CGCGCTCCTAATTATACACAACC | |
| NGS054 | Barley | genetic mapping | CAPS | FORWARD | 61,58 | TTGGAGTGAGGGTTCTGGTAATC | PstI |
| NGS054 | Barley | genetic mapping | CAPS | REVERSE | 61,82 | CTCGACTGCTTCGTCCAGTTTA | |
| NGS084 | Barley | genetic mapping | CAPS | FORWARD | 62,37 | CTTTATTCTCACGTCGTGCACTC | |
| NGS084 | Barley | genetic mapping | CAPS | REVERSE | 60,46 | TGAAGTAGATGCTCCGTCATCCT | BssHII |
| NGS094 | Barley | genetic mapping | CAPS | FORWARD | 61,44 | TGCAAGAGCATCTTCCCTTCTT | HhaI |
| NGS094 | Barley | genetic mapping | CAPS | REVERSE | 60,28 | CTTGCCAACATGCCAAGAGTAG | |
| NGS158 | Barley | genetic mapping | CAPS | FORWARD | 61,71 | TCAACTACACAAGTTCCCGAATTAAC | BsmAI |
| NGS158 | Barley | genetic mapping | CAPS | REVERSE | 62,41 | TGTGAGTCATCAAGGTCCAAGG | |
| NGS160_F | Barley | genetic mapping | CAPS | FORWARD | 62,1 | GTGGCATCATTAGCATAGGATTACTG | HgaI |
| NGS160_R | Barley | genetic mapping | CAPS | REVERSE | 61,72 | AATATGCAAGGTACACCACAAAGAAG | |
| NGS163 | Barley | genetic mapping | CAPS | FORWARD | 62,12 | CGTATCCGGTGTATCGACGTATT | BsrGI |
| NGS163 | Barley | genetic mapping | CAPS | REVERSE | 62,07 | TTATCTTCTCTAGAGTGCTGGCTTGA | |
| NGS164 | Barley | genetic mapping | CAPS | FORWARD | 63,37 | AGCCATGGGCCATTATCTTAATTATC | HhaI |
| NGS164 | Barley | genetic mapping | CAPS | REVERSE | 62,83 | CATGGAATGCACAACTCCTATGTC | |
| NGS166 | Barley | genetic mapping | CAPS | FORW | 61, | AATTCCTGAAACAACGATCAA | Hpy99I |

| | | | | | | | |
|---|---|---|---|---|---|---|---|
| | | | | ARD | 83 | GTTC | |
| NGS166 | Barley | genetic mapping | CAPS | REVE RSE | 62, 28 | CAAGTATGTAATGTTGTGGTGA AGCA | |
| NGS168 | Barley | genetic mapping | CAPS | FORW ARD | 61, 82 | CAATATACGCCGTGTCATACTC TCTT | NdeI |
| NGS168 | Barley | genetic mapping | CAPS | REVE RSE | 62, 27 | ATATGATCGAGTGGACTGGGA GTT | |
| NGS169 | Barley | genetic mapping | CAPS | FORW ARD | 62, 35 | GAGTAGAGGCGCACAGGTGTC | HpyCH4 1V |
| NGS169 | Barley | genetic mapping | CAPS | REVE RSE | 61, 75 | CCAGACATTCTCATTGAAAGA GCTAC | |
| NGS142_F | Barley | genetic mapping | CAPS | FORW ARD | | GACGGCCCTGGTATTAGATAT G | MboII |
| NGS142_R | Barley | genetic mapping | CAPS | REVE RSE | | CCCATCATAACCAAACAGTCCT | |
| TCP-CDS_F2 | Barley | COM1_CDS_resequening | - | FORW ARD | 61, 93 | AGGAAGAAGAGAGTCCTCAAC CAA | |
| TCP-CDS_R2 | Barley | COM1_CDS_resequening | - | REVE RSE | 62, 88 | TAAGCTGCTCGATCGCTAGTAC CT | |
| TCP-CDS_F | Barley | COM1_CDS_resequening | - | FORW ARD | 63, 62 | ATATTGTACTCAAGTGCAGGC AGCTACTA | |
| TCP-CDS_R | Barley | COM1_CDS_resequening | - | REVE RSE | 62, 13 | CATGCAATAATTAACTAAGAA CATGATGC | |
| FPC44150-37B_F | Barley | COM1_promotor region | - | FORW ARD | 61 | CACTGTCTATGGAGAGACCAC ATAGATT | |
| FPC44150-37B_R | Barley | COM1_promotor region | - | REVE RSE | 61 | GTGAGCTAGGCAGCTAGGTAT TTATTAG | |
| FPC44150-38_F | Barley | COM1_promotor region | - | FORW ARD | 62, 34 | TGTTTTCTACTAGTGTCAAGAA CCCTACC | |
| FPC44150-38_R | Barley | COM1_promotor region | - | REVE RSE | 62, 54 | GAGAAAATGTGAGTTATCCTG AACCAG | |
| xNGS129_F | Barley | qPCR | - | FORW ARD | 60 | CGAGCGCATCATGTTCTTAGTT AAT | |
| xNGS129_R | Barley | qPCR | - | REVE | 60 | AGCAACATAGAACAAAACCAT | |

| | | | | | | |
|---|---|---|---|---|---|---|
| | | | | RSE | | GAGAT |
| Com1_ISH_1F20 | Barley | In Situ | - | FORW ARD | | CACAGACGCGAGATGAACAG |
| Com1_ISH_1R20 | Barley | In Situ | - | REVE RSE | | AAAAGGCATCACCCTCAAAA |
| ActinGene | Barley | qPCR | | FORW ARD | | |
| ActinGene | Barley | qPCR | | REVE RSE | | |
| TILLING_Bd-TCP_F1 | Brachyp odium | Screening of TILLING population | - | FORW ARD | 57. 10 | GCAGCAGCAGCAAACTACTA |
| TILLING_Bd-TCP_R1 | Brachyp odium | Screening of TILLING population | - | REVE RSE | 57. 14 | GCTTGGACTGAGTGAGCAG |
| TILLING_Bd-TCP_F2 | Brachyp odium | Screening of TILLING population | - | FORW ARD | 58. 48 | TTTGACAAGGCCAGCAAG |
| TILLING_Bd-TCP_R2 | Brachyp odium | Screening of TILLING population | - | REVE RSE | 58. 25 | AACCACACGCAACAAAGC |
| BdACTIN2_F (Bradi1g10630) | Brachyp odium | qPCR | | FORW ARD | | GTCGTTGCTCCTCCTGAAAG |
| BdACTIN2_R (Bradi1g10630) | Brachyp odium | qPCR | | REVE RSE | | ATCCACATCTGCTGGAAGGT |
| Bd_TCP_InSitu_F | Brachyp odium | qPCR | | FORW ARD | 61. 67 | CAGACCAAGTTCAGCAGAGAT GTAG |
| Bd_TCP_InSitu_R | Brachyp odium | qPCR | | REVE RSE | 61. 72 | CCATCCAAATCAAGAGGTGTA CTTT |
| TILL_SbTCP_F1 | Sorghu m | Screening of TILLING population | | FORW ARD | 62. 25 | GAAGAAGCAGTAGCAGTGGCA GTA |
| TILL_SbTCP_R1 | Sorghu m | Screening of TILLING population | | REVE RSE | 62. 26 | CTTGCTGGCCTTATCGAAGC |
| TILL_SbTCP_F2 | Sorghu m | Screening of TILLING population | | FORW ARD | 62. 11 | ATGCGGTTGTCCCTCGAC |
| TILL_SbTCP_R2 | Sorghu m | Screening of TILLING population | | REVE RSE | 62. 10 | GATAGTGAAGAAGTGCTTGCC AGA |
| TILL_SbTCP_F3 | Sorghu | Screening of TILLING population | | FORW | 62. | CACTACGTGGACCATCACTTCT |

| | | | | | | |
|---|---|---|---|---|---|---|
| | m | | | ARD | 17 | TC |
| TILL_SbTCP_R3 | Sorghum | Screening of TILLING population | | REVERSE | 62.64 | CTAGAGCTCAACTTCTCGGCAACT |
| SbActin_F | Sorghum | qPCR | | FORWARD | | TGGCATCTCTCAGCACATTC |
| SbActin_R | Sorghum | qPCR | | REVERSE | | GGGCGGAAAGAATTAGAAGC |
| Sb_TCP_qPCR_F | Sorghum | qPCR | | FORWARD | 60,07 | CACTAGTAGCTAGCTCTTTCTTTATCTGG |
| Sb_TCP_qPCR_R | Sorghum | qPCR | | REVERSE | 60,42 | CCAGTAGCATTAACTTAAAGGAGTTCA |
| Com2-Bw_SfiI_F | Barley | To screen/sequence for A300C com2 Haplotype in com1.a/com2.g DM population | | FORWARD | 62.51 | CTCCCAGATGATGGCGTTCT |
| Com2-Bw_SfiI_R | Barley | To screen/sequence for A300C com2 Haplotype in com1.a/com2.g DM population | | REVERSE | 62.94 | GAACGGCGGGTAGTTGTTGTAG |
| iLL_TIL_BdTCP_F1 | Brachypodium | illumina sequencing for TILLING mutation detection | | | | TTCCCTACACGACGCTCTTCCGATCTGCAGCAGCAGCAAACTACTA |
| iLL_TIL_BdTCP_R1 | Brachypodium | illumina sequencing for TILLING mutation detection | | | | AGTTCAGACGTGTGCTCTTCCGATCTGCTTGGACTGAGTGAGCAG |
| iLL_TIL_BdTCP_F2 | Brachypodium | illumina sequencing for TILLING mutation detection | | | | TTCCCTACACGACGCTCTTCCGATCTTTTGACAAGGCCAGCAAG |
| iLL_TIL_BdTCP_R2 | Brachypodium | illumina sequencing for TILLING mutation detection | | | | AGTTCAGACGTGTGCTCTTCCGATCTAACCACACGCAACAAAGC |

Supplementary Table 2. Graphical genotyping of the critical F2 recombinants used to develop F3 families

| | Critical F2 recombinants_selected to develop F3 | | | | | | | | | | | | | | |
|---|---|---|---|---|---|---|---|---|---|---|---|---|---|---|---|
| | 1 | 2 | 3 | 4 | 5 | 6 | 7 | 8 | 9 | 10 | 11 | 12 | 13 | 14 | 15 |
| Marker ID | 4839 | 5176 | 5250 | 5403 | 5409 | 5443 | 5504 | 4953 | 5326 | 5483 | 5070 | 5045 | 174 | 5407 | 5488 |
| NGS065 | h | h | h | h | h | h | b | b | b | b | a | a | a | a | a |
| NGS066 | h | h | h | h | h | h | b | b | b | b | a | a | a | a | a |
| NGS046 | h | h | h | h | h | h | b | b | b | b | a | a | a | a | a |
| NGS083 | h | h | h | h | h | h | b | b | b | b | a | a | a | a | a |
| d1652 | h | h | h | h | h | h | b | b | b | b | a | a | a | a | a |
| NGS049 | h | h | h | h | h | h | b | b | b | b | a | a | a | a | a |
| NGS084 | h | h | h | h | h | h | b | b | b | b | a | a | a | a | a |
| NGS168 | h | h | h | h | h | h | b | b | b | b | a | a | a | h | a |
| NGS169 | h | b | b | h | h | h | b | b | h | b | h | h | h | h | a |
| NGS166 | h | b | b | h | h | h | b | b | h | b | h | h | h | h | a |
| *com1.a/Phenotype* | c | c | c | c | c | c | c | c | c | c | c | c | c | c | a |
| NGS164 | h | b | b | h | h | h | b | b | h | b | h | h | h | h | a |
| NGS163 | h | b | b | h | h | h | b | b | h | b | h | h | h | h | a |
| NGS160 | h | b | b | h | h | h | b | b | h | b | h | h | h | h | a |
| NGS158 | b | b | b | h | h | h | b | h | h | b | h | h | h | h | a |
| *NGS094* | b | b | b | b | b | b | h | h | h | h | h | h | h | h | h |
| NGS111 | b | b | b | b | b | b | h | h | h | h | h | h | h | h | h |
| NGS112 | b | b | b | b | b | b | h | h | h | h | h | h | h | h | h |
| *NGS099* | b | b | b | b | b | b | h | h | h | h | h | h | h | h | h |
| *NGS142* | b | b | b | b | b | b | h | h | h | h | h | h | h | h | h |

| Supplementary Table 3. Genotypic and phenotypic data of the F3 progenies | | | | | | |
|---|---|---|---|---|---|---|
| CriticalRecF2_ID | GenotypeParentF2_closely linked marker (NGS164 & NGS166) | CriticalRec_ProgenyF3_ID | Genotype_ProgenyF3_NGS169 | Phenotype | Number of Spike | Number of Branched Spike |
| 174 | h | 174 - 1 | h | Wt | 16 | - |
| 174 | h | 174 - 2 | h | Wt | 18 | - |
| 174 | h | 174 - 3 | h | Wt | 17 | - |
| 174 | h | 174 - 4 | a | branched | 17 | 13 |
| 174 | h | 174 - 5 | b | Wt | 20 | - |
| 174 | h | 174 - 6 | h | Wt | 17 | - |
| 174 | h | 174 - 7 | b | Wt | 22 | - |
| 174 | h | 174 - 8 | b | Wt | 21 | - |
| 174 | h | 174 - 9 | a | branched | 15 | 12 |
| 174 | h | 174 - 10 | a | branched | 19 | 16 |
| 174 | h | 174 - 12 | a | branched | 14 | 8 |
| 174 | h | 174 - 13 | h | Wt | 12 | - |
| 174 | h | 174 - 14 | h | Wt | 16 | - |
| 174 | h | 174 - 15 | h | Wt | 14 | - |
| 174 | h | 174 - 16 | b | Wt | 17 | - |
| 4839 | h | 4839 - 1 | h | Wt | 15 | - |
| 4839 | h | 4839 - 2 | b | Wt | 15 | - |
| 4839 | h | 4839 - 3 | a | branched | 15 | 6 |
| 4839 | h | 4839 - 4 | b | Wt | 15 | - |
| 4839 | h | 4839 - 5 | a | branched | 15 | 5 |
| 4839 | h | 4839 - 6 | b | Wt | 13 | - |
| 4839 | h | 4839 - 7 | h | Wt | 15 | - |
| 4839 | h | 4839 - 8 | h | Wt | 15 | - |

| | | | | | | |
|---|---|---|---|---|---|---|
| 4839 | h | 4839 - 9 | a | branched | 20 | 12 |
| 4839 | h | 4839 - 10 | b | Wt | 14 | - |
| 4839 | h | 4839 - 11 | h | Wt | 16 | - |
| 4839 | h | 4839 - 12 | h | Wt | 19 | - |
| 4839 | h | 4839 - 13 | h | Wt | 20 | - |
| 4839 | h | 4839 - 14 | h | Wt | 19 | - |
| 4839 | h | 4839 - 15 | h | Wt | 19 | - |
| 4839 | h | 4839 - 16 | h | Wt | 12 | - |
| 4953 | b | 4953 - 1 | b | Wt | 13 | - |
| 4953 | b | 4953 - 2 | b | Wt | 15 | - |
| 4953 | b | 4953 - 3 | b | Wt | 14 | - |
| 4953 | b | 4953 - 4 | b | Wt | 13 | - |
| 4953 | b | 4953 - 5 | b | Wt | 16 | - |
| 4953 | b | 4953 - 6 | b | Wt | 17 | - |
| 4953 | b | 4953 - 7 | b | Wt | 13 | - |
| 4953 | b | 4953 - 8 | b | Wt | 18 | - |
| 4953 | b | 4953 - 9 | b | Wt | 13 | - |
| 4953 | b | 4953 - 10 | b | Wt | 15 | - |
| 4953 | b | 4953 - 11 | b | Wt | 19 | - |
| 4953 | b | 4953 - 12 | b | Wt | 16 | - |
| 4953 | b | 4953 - 13 | b | Wt | 19 | - |
| 4953 | b | 4953 - 14 | b | Wt | 14 | - |
| 4953 | b | 4953 - 15 | b | Wt | 15 | - |
| 4953 | b | 4953 - 16 | b | Wt | 17 | - |
| 5045 | h | 5045 - 1 | a | branched | 19 | 10 |
| 5045 | h | 5045 - 2 | h | Wt | 14 | - |
| 5045 | h | 5045 - 3 | h | Wt | 11 | - |
| 5045 | h | 5045 - 5 | h | Wt | 14 | - |
| 5045 | h | 5045 - 6 | h | Wt | 16 | - |

| | | | | | | |
|---|---|---|---|---|---|---|
| 5045 | h | 5045 - 7 | h | Wt | 20 | - |
| 5045 | h | 5045 - 8 | a | branched | 15 | 6 |
| 5045 | h | 5045 - 9 | h | Wt | 15 | - |
| 5045 | h | 5045 - 10 | h | Wt | 18 | - |
| 5045 | h | 5045 - 11 | a | branched | 12 | 5 |
| 5045 | h | 5045 - 13 | h | Wt | 20 | - |
| 5045 | h | 5045 - 14 | h | Wt | 19 | - |
| 5045 | h | 5045 - 15 | h | Wt | 15 | - |
| 5045 | h | 5045 - 16 | h | Wt | 15 | - |
| 5070 | h | 5070 - 1 | h | Wt | 16 | - |
| 5070 | h | 5070 - 2 | h | Wt | 17 | - |
| 5070 | h | 5070 - 3 | h | Wt | 15 | - |
| 5070 | h | 5070 - 4 | a | branched | 16 | 11 |
| 5070 | h | 5070 - 5 | b | Wt | 20 | - |
| 5070 | h | 5070 - 6 | h | Wt | 21 | - |
| 5070 | h | 5070 - 7 | a | branched | 17 | 9 |
| 5070 | h | 5070 - 8 | a | branched | 16 | 8 |
| 5070 | h | 5070 - 9 | b | Wt | 18 | - |
| 5070 | h | 5070 - 11 | b | Wt | 25 | - |
| 5070 | h | 5070 - 12 | a | branched | 14 | 9 |
| 5070 | h | 5070 - 13 | h | Wt | 15 | - |
| 5070 | h | 5070 - 14 | h | Wt | 14 | - |
| 5070 | h | 5070 - 15 | h | Wt | 26 | - |
| 5070 | h | 5070 - 16 | a | branched | 19 | 11 |
| 5176 | b | 5176 - 1 | b | Wt | 13 | - |
| 5176 | b | 5176 - 2 | b | Wt | 16 | - |
| 5176 | b | 5176 - 3 | b | Wt | 21 | - |
| 5176 | b | 5176 - 4 | b | Wt | 13 | - |
| 5176 | b | 5176 - 5 | b | Wt | 14 | - |

| | | | | | | |
|---|---|---|---|---|---|---|
| 5176 | b | 5176 - 6 | b | Wt | 16 | - |
| 5176 | b | 5176 - 7 | b | Wt | 28 | - |
| 5176 | b | 5176 - 8 | b | Wt | 12 | - |
| 5176 | b | 5176 - 9 | b | Wt | 14 | - |
| 5176 | b | 5176 - 10 | b | Wt | 25 | - |
| 5176 | b | 5176 - 11 | b | Wt | 16 | - |
| 5176 | b | 5176 - 12 | b | Wt | 12 | - |
| 5176 | b | 5176 - 13 | b | Wt | 13 | - |
| 5176 | b | 5176 - 14 | b | Wt | 13 | - |
| 5176 | b | 5176 - 15 | b | Wt | 19 | - |
| 5176 | b | 5176 - 16 | b | Wt | 16 | - |
| 5250 | b | 5250 - 1 | b | Wt | 13 | - |
| 5250 | b | 5250 - 2 | b | Wt | 13 | - |
| 5250 | b | 5250 - 3 | b | Wt | 18 | - |
| 5250 | b | 5250 - 4 | b | Wt | 16 | - |
| 5250 | b | 5250 - 5 | b | Wt | 13 | - |
| 5250 | b | 5250 - 6 | b | Wt | 12 | - |
| 5250 | b | 5250 - 7 | b | Wt | 14 | - |
| 5250 | b | 5250 - 8 | b | Wt | 17 | - |
| 5250 | b | 5250 - 9 | b | Wt | 16 | - |
| 5250 | b | 5250 - 10 | b | Wt | 13 | - |
| 5250 | b | 5250 - 11 | b | Wt | 13 | - |
| 5250 | b | 5250 - 12 | b | Wt | 15 | - |
| 5250 | b | 5250 - 13 | b | Wt | 21 | - |
| 5250 | b | 5250 - 14 | b | Wt | 18 | - |
| 5250 | b | 5250 - 15 | b | Wt | 15 | - |
| 5250 | b | 5250 - 16 | b | Wt | 16 | - |
| 5326 | h | 5326 - 1 | h | Wt | 16 | - |
| 5326 | h | 5326 - 2 | b | Wt | 17 | - |

| | | | | | | |
|---|---|---|---|---|---|---|
| 5326 | h | 5326 - 3 | h | Wt | 15 | - |
| 5326 | h | 5326 - 4 | h | Wt | 15 | - |
| 5326 | h | 5326 - 5 | b | Wt | 13 | - |
| 5326 | h | 5326 - 6 | b | Wt | 16 | - |
| 5326 | h | 5326 - 7 | h | Wt | 17 | - |
| 5326 | h | 5326 - 8 | h | Wt | 16 | - |
| 5326 | h | 5326 - 9 | a | branched | 16 | 8 |
| 5326 | h | 5326 - 10 | b | Wt | 11 | - |
| 5326 | h | 5326 - 11 | b | Wt | 15 | - |
| 5326 | h | 5326 - 12 | h | Wt | 16 | - |
| 5326 | h | 5326 - 13 | a | branched | 13 | 10 |
| 5326 | h | 5326 - 14 | b | Wt | 12 | - |
| 5326 | h | 5326 - 15 | b | Wt | 11 | - |
| 5326 | h | 5326 - 16 | h | Wt | 15 | - |
| 5403 | h | 5403 - 1 | h | Wt | 15 | - |
| 5403 | h | 5403 - 2 | h | Wt | 16 | - |
| 5403 | h | 5403 - 3 | a | branched | 17 | 13 |
| 5403 | h | 5403 - 4 | h | Wt | 13 | - |
| 5403 | h | 5403 - 5 | b | Wt | 17 | - |
| 5403 | h | 5403 - 6 | a | branched | 14 | 9 |
| 5403 | h | 5403 - 7 | a | branched | 16 | 10 |
| 5403 | h | 5403 - 8 | h | Wt | 15 | - |
| 5403 | h | 5403 - 9 | h | Wt | 13 | - |
| 5403 | h | 5403 - 10 | b | Wt | 16 | - |
| 5403 | h | 5403 - 12 | b | Wt | 18 | - |
| 5403 | h | 5403 - 13 | h | Wt | 15 | - |
| 5403 | h | 5403 - 14 | b | Wt | 15 | - |
| 5403 | h | 5403 - 15 | a | branched | 15 | 9 |
| 5403 | h | 5403 - 16 | b | Wt | 16 | - |

| | | | | | | |
|---|---|---|---|---|---|---|
| 5407 | h | 5407 - 1 | a | branched | 15 | 8 |
| 5407 | h | 5407 - 2 | h | Wt | 19 | - |
| 5407 | h | 5407 - 3 | b | Wt | 13 | - |
| 5407 | h | 5407 - 4 | h | Wt | 14 | - |
| 5407 | h | 5407 - 5 | h | Wt | 14 | - |
| 5407 | h | 5407 - 6 | a | branched | 15 | 11 |
| 5407 | h | 5407 - 7 | h | Wt | 14 | - |
| 5407 | h | 5407 - 8 | h | Wt | 15 | - |
| 5407 | h | 5407 - 9 | h | Wt | 15 | - |
| 5407 | h | 5407 - 10 | b | Wt | 13 | - |
| 5407 | h | 5407 - 11 | h | Wt | 16 | - |
| 5407 | h | 5407 - 12 | a | branched | 18 | 10 |
| 5407 | h | 5407 - 13 | a | branched | 16 | 12 |
| 5407 | h | 5407 - 14 | h | Wt | 15 | - |
| 5407 | h | 5407 - 15 | h | Wt | 14 | - |
| 5407 | h | 5407 - 16 | h | Wt | 15 | - |
| 5409 | h | 5409 - 1 | - | Wt | 16 | - |
| 5409 | h | 5409 - 2 | h | Wt | 13 | - |
| 5409 | h | 5409 - 3 | h | Wt | 13 | - |
| 5409 | h | 5409 - 4 | b | Wt | 13 | - |
| 5409 | h | 5409 - 5 | h | Wt | 14 | - |
| 5409 | h | 5409 - 6 | a | branched | 15 | 11 |
| 5409 | h | 5409 - 7 | a | branched | 19 | 7 |
| 5409 | h | 5409 - 8 | h | Wt | 13 | - |
| 5409 | h | 5409 - 9 | h | Wt | 16 | - |
| 5409 | h | 5409 - 11 | h | Wt | 19 | - |
| 5409 | h | 5409 - 13 | h | Wt | 13 | - |
| 5409 | h | 5409 - 14 | a | branched | 16 | 11 |
| 5409 | h | 5409 - 15 | b | Wt | 13 | - |

| | | | | | | |
|---|---|---|---|---|---|---|
| 5409 | h | 5409 - 16 | b | Wt | 14 | - |
| 5443 | h | 5443 - 1 | h | Wt | 14 | - |
| 5443 | h | 5443 - 2 | a | branched | 12 | 7 |
| 5443 | h | 5443 - 3 | h | Wt | 14 | - |
| 5443 | h | 5443 - 4 | b | Wt | 13 | - |
| 5443 | h | 5443 - 5 | h | Wt | 13 | - |
| 5443 | h | 5443 - 6 | h | Wt | 18 | - |
| 5443 | h | 5443 - 7 | a | branched | 17 | 10 |
| 5443 | h | 5443 - 8 | a | branched | 12 | 9 |
| 5443 | h | 5443 - 9 | b | Wt | 14 | - |
| 5443 | h | 5443 - 11 | h | Wt | 12 | - |
| 5443 | h | 5443 - 12 | h | Wt | 14 | - |
| 5443 | h | 5443 - 13 | h | Wt | 14 | - |
| 5443 | h | 5443 - 14 | a | branched | 16 | 12 |
| 5443 | h | 5443 - 15 | a | branched | 14 | 10 |
| 5443 | h | 5443 - 16 | b | Wt | 15 | - |
| 5483 | b | 5483 - 1 | b | Wt | 15 | - |
| 5483 | b | 5483 - 2 | b | Wt | 15 | - |
| 5483 | b | 5483 - 3 | b | Wt | 15 | - |
| 5483 | b | 5483 - 4 | b | Wt | 13 | - |
| 5483 | b | 5483 - 5 | b | Wt | 11 | - |
| 5483 | b | 5483 - 6 | b | Wt | 13 | - |
| 5483 | b | 5483 - 7 | b | Wt | 16 | - |
| 5483 | b | 5483 - 8 | b | Wt | 14 | - |
| 5483 | b | 5483 - 9 | b | Wt | 15 | - |
| 5483 | b | 5483 - 10 | b | Wt | 14 | - |
| 5483 | b | 5483 - 11 | b | Wt | 14 | - |
| 5483 | b | 5483 - 12 | b | Wt | 16 | - |
| 5483 | b | 5483 - 13 | b | Wt | 12 | - |

| | | | | | | |
|---|---|---|---|---|---|---|
| 5483 | b | 5483 - 14 | b | Wt | 14 | - |
| 5483 | b | 5483 - 15 | b | Wt | 15 | - |
| 5483 | b | 5483 - 16 | b | Wt | 13 | - |
| 5488 | a | 5488 - 2 | a | branched | 22 | 15 |
| 5488 | a | 5488 - 3 | a | branched | 23 | 14 |
| 5488 | a | 5488 - 5 | a | branched | 19 | 14 |
| 5488 | a | 5488 - 6 | a | branched | 12 | 10 |
| 5488 | a | 5488 - 7 | a | branched | 21 | 15 |
| 5488 | a | 5488 - 8 | a | branched | 17 | 13 |
| 5488 | a | 5488 - 9 | a | branched | 20 | 11 |
| 5488 | a | 5488 - 10 | a | branched | 20 | 10 |
| 5488 | a | 5488 - 11 | a | branched | 23 | 13 |
| 5488 | a | 5488 - 12 | a | branched | 25 | 16 |
| 5488 | a | 5488 - 13 | a | branched | 20 | 14 |
| 5488 | a | 5488 - 14 | a | branched | 20 | 12 |
| 5488 | a | 5488 - 15 | a | branched | 21 | 12 |
| 5488 | a | 5488 - 16 | a | branched | 20 | 10 |
| 5504 | b | 5504 - 1 | b | Wt | 15 | - |
| 5504 | b | 5504 - 2 | b | Wt | 19 | - |
| 5504 | b | 5504 - 3 | b | Wt | 24 | - |
| 5504 | b | 5504 - 4 | b | Wt | 21 | - |
| 5504 | b | 5504 - 5 | b | Wt | 21 | - |
| 5504 | b | 5504 - 6 | b | Wt | 17 | - |
| 5504 | b | 5504 - 7 | b | Wt | 19 | - |
| 5504 | b | 5504 - 8 | b | Wt | 11 | - |

| Supplementary Table 4. Part 1. List of the TILLING as well as induced mutants per corresponding species | | | | | | | | | | |
|---|---|---|---|---|---|---|---|---|---|---|
| Species | TILLING Line ID | DNA Position (from Start Codon) | SNP type | Homo/ Hetero | Wild aa | Mutant aa | Location within the gene | the grass sp. with deviated aa | SIFT prediction | SIFT score |
| Barley | M3.15104 | 305 | G-to-A | homo | R | K | within TCP Domain | no change in grass; R | NA | NA |
| Barley | M3.4406 | 334 | G-to-A | homo | A | T | within TCP Domain | no change in grass; A | NA | NA |
| Barley | M3.2598 | 346 | C-to-T | homo | R | W | within TCP Domain | no change in grass; R | NA | NA |
| Barley | M3.13729 | 335 | C-to-T | homo | A | V | within TCP Domain | no change in grass; A | NA | NA |
| Barley | M3.4063 | 211 | C-to-T | hetro | P | S | befor TCP Domain | OS: D | NA | NA |
| Barley | M3.9299 | 595 | G-to-A | hetro | G | R | after TCP Domain | no change in grass; G | NA | NA |
| Barley | M3.14287 | 109 | G-to-A | hetro | D | N | befor TCP Domain | OS: Q | NA | NA |
| Barley | M3.2821 | 241 | C-to-T | hetro | p | S | befor TCP Domain | in OS, Tu, BD and SB: A | NA | NA |
| Barley | M3.14325 | 242 | C-to-A | hetro | p | Q | befor TCP Domain | in OS, Tu, BD and SB: A | NA | NA |
| Barley | M3.6995 | 290 | G-to-A | homo | R | K | befor TCP Domain | no change in grass; R | NA | NA |
| Barley | M3.12123 | 290 | G-to-A | hetro | R | K | befor TCP Domain | no change in grass; R | NA | NA |
| Barley | M3.13483 | 290 | G-to-A | hetro | R | K | befor TCP Domain | no change in grass; R | NA | NA |
| Barley | M3.2927 | 496 | C-to-T | homo | L | F | after TCP Domain | OS: S, ZM: E, Sb: L | NA | NA |
| Barley | M3.13996 | 511 | G-to-A | hetro | A | T | after TCP Domain | ZM: R, Sb: V | NA | NA |
| Barley | M3.14152 | 514 | G-to-A | homo | G | R | after TCP Domain | OS: A, ZM: - | NA | NA |
| Barley | M3.6504 | 515 | G-to-A | hetro | G | E | after TCP Domain | OS: A, ZM: - | NA | NA |
| Barley | M3.3717 | 528 | G-to-A | hetro | M | I | after TCP Domain | in OS: R, BD: -, SB: - , ZM: - | NA | NA |
| Barley | M3.11298 | 544 | G-to-A | hetro | V | M | after TCP Domain | in OS: A, BD: -, SB: D , ZM: D | NA | NA |
| Barley | M3.11933 | 544 | G-to-A | homo | V | M | after TCP Domain | in OS: A, BD: -, SB: D , ZM: D | NA | NA |
| Barley | M3.13403 | 544 | G-to-A | hetro | V | M | after TCP Domain | in OS: A, BD: -, SB: D , ZM: D | NA | NA |
| Barley | M3.6448 | 573 | G-to-T | hetro | E | D | after TCP Domain | in OS: V, SB: S , ZM: S | NA | NA |
| Barley | M3.12394 | 595 | G-to-A | hetro | G | R | after TCP Domain | no change in grass; G | NA | NA |

| | | | | | | | | | | |
|---|---|---|---|---|---|---|---|---|---|---|
| Barley | M3.3609 | 635 | G-to-A | homo | R | k | after TCP Domain | no change in grass; R | NA | NA |
| Barley | M3.9852 | 641 | G-to-A | hetro | G | D | after TCP Domain | no change in grass; G | NA | NA |
| Barley | M3.11856 | 670 | C-to-T | homo | R | C | after TCP Domain | in OS: -, BD: L, SB: - , ZM: - | NA | NA |
| Sorghum | ARS152 | 188 | G-to-A | hetro | R | H | Befor TCP Domain | not conserved | NA | NA |
| Sorghum | ARS180 | 430 | A-to-G | homo | A | T | within TCP Domain | no change in grass; A | NA | NA |
| Sorghum | ARS137 | 577 | C-to-T | hetro | P | S | after TCP Domain | not conserved | NA | NA |
| Brachypo dium | 8004 | 173 | C_To_T | | S58F | | Befor TCP Domain | | Damaging | 0 |
| Brachypo dium | 8231 | 187 | G_To_ A | | A63 T | | Befor TCP Domain | | Damaging | 0 |
| Brachypo dium | 8566 | 266 | G_To_ A | | G89 D | | Befor TCP Domain | | Damaging | 0,01 |
| Brachypo dium | 7576 | 278 | C_To_T | | A93 V | | Befor TCP Domain | | Damaging | 0,01 |
| Brachypo dium | 4957 | 295 | C_To_T | | P99S | | Befor TCP Domain | | Damaging | 0,05 |
| Brachypo dium | 5446 | 346 | C_To_T | | Q116 * | | within TCP Domain | | - | - |
| Brachypo dium | 8373 | 437 | G_To_ A | | S146 N | | within TCP Domain | | Damaging | 0,02 |
| Brachypo dium | 5149 | 472 | C_To_T | | P158 S | | within TCP Domain | | Tolerated | 0,07 |
| Brachypo dium | 5337 | 533 | C_To_T | | A178 V | | within TCP Domain | | Tolerated | 0,15 |
| Brachypo dium | 6339 | 536 | G_To_ A | | G179 E | | within TCP Domain | | Tolerated | 0,07 |
| Brachypo dium | 4196 | 785 | G_To_ A | | G262 D | | after TCP Domain | | Damaging | 0 |

Supplementary Table 4. Part 2.

| Running Number | Branched (compositum) Barleys: | Gene Bank | Type | ID | type of mutation and the doner line | Cross with com1a | PCR with TCP_potential promotor based primers | TCP_CDS_Resequening |
|---|---|---|---|---|---|---|---|---|
| 4 | Mut.3906 | GER | Spring | MHOR347 | X-ray; spike with short branches; 1949- 'Heines Haisa'; | Done/allelic | Amplicon observed | No Amplicon |
| 8 | com1.j | USA | Spring | NGB22020 | X-ray; induced mutant in Donaria (PI 161974) isolated by F. Scholz | | No Amplicon | No Amplicon |
| int-h 42 | int-h 42 | SWE | #N/A | #N/A | /neutrons/ KRISTINA | | No Amplicon | No Amplicon |
| int-h 43 | int-h 43 | SWE | #N/A | #N/A | /hydroxy n-propyl methanesulfonate/ KRISTINA | | No Amplicon | No Amplicon |
| int-h 44 | int-h 44 | SWE | #N/A | #N/A | /ethyl methanesulfonate/ KRISTINA | | No Amplicon | No Amplicon |

Supplementary Table 5. Phenotypic data of the sorghum-TILLING mutants

| | | | M5 generation _to measure branch number | | | M6 generation _to measure branch angle | | | | |
|---|---|---|---|---|---|---|---|---|---|---|
| TILLING Line ID_M4 | DNA Position (from Start Codon) | Location within the gene | Number of plants _all homozygous | Average No of nodes/panicle/ family | Average No of branch per nodes/panicle/ family | Number of plants _ all homozygous | No of nodes/panicle/family | Average No of angle per nodes/panicle/ family | Average angle size (degree) per nodes/panicle/ family | Grain formation |
| ARS180 | A430G | within TCP Domain | 8 | 12 | 4,2* | 7 | 3 basal | 10,1 | 5,2*** | only 20-40 grains/panicle |
| ARS137 | C577T | after TCP Domain | 8 | 11,5 | 4,4* | 6 | 3 basal | 9,3 | 15,9 | Complete fertility |
| Wt BT623 | - | - | 7 | 10 | 5,4 | 5 | 3 basal | 8,1 | 10,95 | Complete fertility |

* ; ttest ($P < 0.05$)
*** ; ttest ($P < 0.001$)

#### Supplementary source data

**Supplementary source data 1. Differentially expressed genes in Wt and barley mutant with the corresponding statistical values**
